# ILT7 activation and plasmacytoid dendritic cell response are governed by BST2 determinants that are structurally-distinct

**DOI:** 10.1101/557116

**Authors:** Mariana G. Bego, Nolwenn Miguet, Alexandre Laliberté, Nicolas Aschman, Francine Gerard, Angelique A. Merakos, Winfried Weissenhorn, Éric A. Cohen

## Abstract

The interaction of the human pDC receptor ILT7 with its ligand, BST2, significantly regulates pDC’s TLR-induced innate immune responses. This interaction has critical biological consequences, yet, its structural requirements are not fully characterized. The BST2 ectodomain can be divided in structurally and functionally distinct regions; while the coiled-coil region contains a newly-defined ILT7 binding surface, the N-terminal region appears to negatively modulate ILT7 activation. A stable BST2 homodimer binds to ILT7 but post-binding events associated to the unique BST2 coiled-coil plasticity are required to trigger receptor signaling. Hence, BST2 with an unstable or with a rigid coiled-coil fails to activate ILT7, whereas mutations in the N-terminal region enhance activation. Importantly, the biological relevance of these newly defined domains of BST2 is underscored by the identification of mutations with opposed potential to activate ILT7, which are selected under pathological malignant conditions.

## Introduction

Type I interferons (IFN-I) are key soluble antiviral molecules that are predominantly produced by activated plasmacytoid dendritic cells (pDCs)^1^. Their functions reach far beyond their established role during antimicrobial defense as IFN-I are also linked with regulation of immune cell differentiation, survival and homeostasis as well as control of the cell cycle^2–4^. Hence, IFN-I play an important role in orchestrating the natural immune response to cancer and have inhibitory functions that prevent malignant cellular transformation (reviewed in Zitvogel, et al^5^). However, their biological activities can also have deleterious impacts on surrounding healthy cells. Prolonged IFN-I signaling is associated to excessive inflammation and immune dysfunction^6^ and high levels of IFN-I contributes to aberrant immune activation and development of autoimmune diseases^7^. Furthermore, IFN-Is can also act as a double-edged sword when fighting malignant invasive tumors, with the potential to deploy opposite anti- and pro-tumorigenic outcomes given their direct impact on tumor cells and potentially improper activity on tumor infiltrating immune cells^8^. Thus, IFN-I production and signaling need to be tightly regulated to achieve protective immunity during pathological conditions while avoiding harmful toxicity caused by improper or prolonged IFN signaling.

One way to control IFN-I production involves the engagement of pDC-specific regulatory receptors BDCA-2 (CD303) and ILT7 (LILRA4, CD85g). Crosslinking of either regulatory receptor efficiently suppresses the production of IFN-I and other cytokines in response to toll-like receptor 7 and 9 (TLR7/9) activation^9,10^. Interestingly, the natural ligand of ILT7 was found to be BST2 (bone-marrow stromal cell antigen 2), a membrane-associated protein that is itself induced by IFN-I^11^. Given the IFN-I-inducible nature of the ILT7 ligand, it was proposed that BST2 contributes to a negative feedback mechanism controlling IFN-I overproduction by pDCs after viral infection and/or sustained inflammatory responses^11–14^. Remarkably, BST2 expression is constitutively elevated in various cancers such as myelomas, lung cancer, breast cancer, colorectal cancer, and pancreatic cancer^15^. Indeed, constitutive expression of BST2 by human breast cancer cell line and melanoma lines was shown to suppress IFN-I production by pDC via ILT7, raising the possibility that the interaction of BST2 with ILT7 might be contributing to tumor immune suppression and pDC-tumor crosstalk^14^.

BST2 is a small, evolutionary conserved, single-pass type II membrane protein. It has a unique topology as its ectodomain is anchored to the plasma membrane via a N-terminal transmembrane domain and a C-terminal glycophosphatidylinositol (GPI) anchor^16^ (Fig. 1A). BST2 contains a short cytoplasmic tail involved in NF-κB signaling and AP-2-dependent endocytosis^17,18^. The protein is heavily glycosylated on two conserved extracellular asparagine residues (N65 and N92) and forms a stable disulfide-linked dimer via any of the three conserved cysteine residues located towards the N-terminal region of the ectodomain (C53, C63 and C91)^16,19–22^.

**Fig 1.**
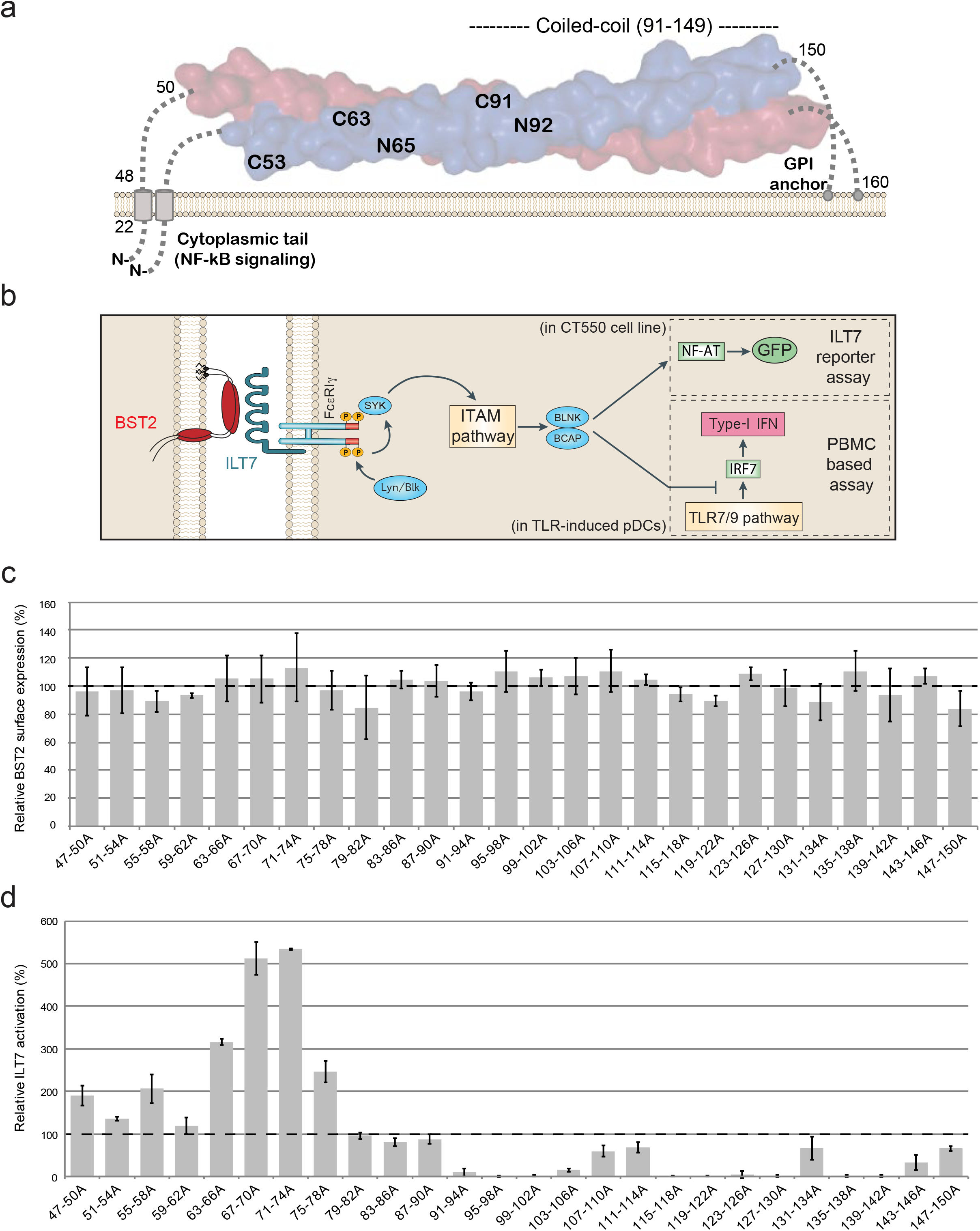
Uncovering BST2 regions relevant for ILT7 activation using an alanine scan analysis of its ectodomain. **a,** Schematic representation of BST2, a type II transmembrane (TM) protein of 160 amino acids, overlaying the crystal structure of BST2 ectodomain (PDB ID: 2XG7) published by Schubert et al ^24^ generated using NGL viewer^51^. BST2 features a short cytoplasmic N-terminus containing di-phosphotyrosines required for NF-κB signaling followed by an α-helical single-pass TM domain and an ectodomain comprising an extended coiled-coil linked back to the plasma membrane by a C-terminal GPI anchor. N-glycosylation sites (N65 and N92) as well as cysteine residues used for disulfide-bond formation (C53, C63 and C91) in the extracellular domain are indicated. **b,** Schematic representation of BST2-ILT7 activation pathways and of the two assays used to measure BST2-mediated ILT7 activation. For the ILT7 reporter assay, BST2-expressing HEK-293T cells are co-cultured with ILT7+ NFAT-GFP reporter cells for 18-24 h and activation of the ITAM pathways is measured as the percentage of GFP+ reporter cells by flow cytometry. For the PBMC-based assay, BST2-expressing HEK-293T cells are co-cultured with PBMCs. After 4 h of co-culture, samples are either untreated or treated with Gardiquimod (TLR7 agonist) and levels of bioactive IFN-I released in supernatants measured 18-24 h later, as described in the Methods section. **c-d,** Alanine scan of BST2 ectodomain (non-overlapping groups of 4 residues mutated to alanines from positions 47 to 150). **c,** Relative BST2 surface expression in HEK-293T cells transfected with empty plasmid, plasmid encoding for BST2 WT or alanine mutants (n=6). Percentages of mean fluorescence intensities (MFIs) were calculated relative to BST2 WT-expressing cells (100%). **d,** ILT7+ NFAT-GFP reporter cells were co-cultured with control (empty) or HEK-293T cells expressing the above-mentioned BST2 (WT or mutants) and analyzed by flow cytometry (n=6). Percentage of ILT7 activation was plotted as % of GFP+ cells in each condition relative to the BST2 WT condition (100%) after subtracting the % of GFP+ cells in the no BST2 condition (0%). Error bars represent standard deviation (SD).

The stucture of the human and mouse BST2 ectodomain dimers have been solved by X-ray crystallography. Each monomer consists of a continuous alpha helix organized in parallel orientation, forming a homodimer with a coiled-coil structure stabilized by intermolecular disulfide linkages^23–26^. Biophysical studies indicate that the recombinant coiled-coil BST2 ectodomain dimer exists as a rod-like structure of 15 to 17nm in length with a slight bend at about one third from its N-terminus^26,27^. The ectodomain dimer is hinged at two positions (A88 and G109 in the human protein), giving the N-terminus a degree of rotational flexibility^24–26^. In both species, the ectodomain two-third C-terminal region is arranged as a parallel dimeric coiled-coil formed by 6–7 heptad repeats starting at position C91^23–26^. The presence of a number of destabilizing residues in key central heptad positions confer the ectodomain coiled-coil its characteristic dynamic instability that requires intermolecular disulfides for stability^23^. Such an evolutionary conserved design was proposed to provide the BST2 dimer an inherent plasticity that allows the protein to adapt during dynamic events^23,27^.

BST2 is a multifaceted protein. It is also referred as ‘tetherin’, as it acts as a broad antiviral restriction factor through its ability to physically retain enveloped viruses at the cell surface, thus preventing their release from infected cells^28,29^. We recently reported that while ‘tethering’ human immunodeficiency virus type 1 (HIV-1), BST2 is unable to interact with ILT7 and block IFN-I production by pDCs^30^. Remarkably, the pandemic HIV-1 group M has evolved mechanisms to maintain BST2-ILT7 interaction and limit pDC activation despite its ability to effectively counteract BST2 for efficient HIV-1 release^30^. This property, which limits pDC antiviral responses, was not conserved in the endemic HIV-1 group O^30,31^. While the domains and structural features of BST2 required for tethering HIV-1 have been precisely defined^22,23,28,32,33^, the determinants required for engaging and activating ILT7 are largely undefined. Understanding the structural features of BST2 that are required to engage and activate ILT7 may guide the development of new therapeutic options aimed at restoring pDC functionality and triggering IFN-I production to induce effective antitumor or antiviral immunity.

Using functional assays measuring ILT7 activation by BST2 as well as biophysical studies of BST2 interaction with ILT7, we show that the structurally distinct N- and C-terminal regions of BST2 ectodomain have discrete functions during ILT7 activation. While the unique and highly conserved C-terminal coiled-coil region of BST2 contains key residues required for ILT7 binding, the N-terminal and its central connecting flexible region acts as a modulatory region negatively regulating ILT7 activation. Indeed, we provide evidence that after the initial interaction, the coiled-coil internal flexibility is required for ILT7 activation suggesting a post-binding conformational rearrangement of the BST2/ILT7complex. We further demonstrate that a stable BST2 homodimer is required to trigger ILT7 activation and that both BST2 monomers need to have intact ILT7-binding surface for this function. Importantly, analysis of somatic mutations identified in particular tumor tissues and mapping to these newly defined BST2 functional domains reveals that specific genetic changes in BST2 are capable of either greatly enhancing ILT7-mediated suppression of IFN-I production or completely abrogating this phenotype, thus underscoring the clinical relevance of our findings.

## Results

### The highly conserved coiled-coil region of BST2 is required for ILT7 activation

We previously demonstrated that the BST2 ectodomain is sufficient to interact with ILT7 and activate its inhibitory signaling cascade^30^. To identify the determinants in BST2 ectodomain required for ILT7 activation we used two previously described co-culture assays, namely a reconstituted ILT7 reporter assay and a PBMC-based assay (Fig. 1B). The first functional assay relies on a mouse NFAT-GFP reporter cell line, which also expresses human ILT7 and the mouse FcϵRIγ chain (CT550) such that levels of GFP expression directly correlates with the degree of BST2-mediated ILT7 activation^10,11^. Selected BST2 mutants were also tested in a PBMC-based co-culture assay, which directly measures the quantity of IFN-I produced by PBMCs after TLR7 agonist stimulation following engagement and activation of the ILT7 pathway in pDCs by BST2^30,34^. Given that the extent of ILT7 activation is directly proportional to the surface levels of BST2 expressed on co-cultured 293T targets cells^30^, expression of all BST2 mutants was standardized to match that of the internal BST2 WT control (Fig. S1). A series of alanine substitution mutants starting at amino acid position 47 and continuing through position 150^32^, where each mutant in the panel incorporates four consecutive alanine residues, was used to identify determinants required for ILT7 activation. HEK-293T cells expressing BST2 (WT or mutants) were co-cultured with ILT7^+^ NFAT-GFP reporter cells prior to flow cytometry analysis. The alanine scan analysis revealed two discrete domains in BST2 that modulate ILT7 activation. A first region spanning residues 47 to 90 appeared to act as a modulatory domain as alanine substitutions enhanced BST2-mediated ILT7 activation. In contrast, a second region overlapping the BST2 coiled-coil domain from residues 91 to 147 seemed to have a direct role in ILT7 activation as many alanine mutations resulted in a strong impairment of the ILT7 activation phenotype (Fig. 1C-D).

### A surface required for ILT7 activation is defined by two spatially adjacent residues within the BST2 coiled-coil

We confirmed that the BST2 coiled-coil region was sufficient to interact with ILT7 by surface plasmon resonance *in vitro* (K_D_=2.62 μM) using recombinant GST-tagged truncated BST2 (residues 80 to 147) and baculovirus-expressed soluble ILT7 (Fig. 2A). The primary structure of the coiled-coils is characterized by a periodicity of seven residues or heptad repeat pattern, which are usually labeled *abcdefg*. Heptad positions *a* and *d* (typically hydrophobic residues) form the core present at the interface of the two helices, while *e* and *g* positions (typically charged residues) form inter-helical ionic interactions and are all involved in dimer stabilization^35,36^. Some of the alanine residues introduced during the alanine scan could potentially disrupt the stability of the coiled-coil structure, especially those that substitute amino acid residues located at the center of the α–helix (heptad positions *a* and *d)* or those involved in stabilizing inter-helical bonds (heptad positions *e* and *g*; Fig. 2B-C). Furthermore, as the BST2 ectodomain parallel dimer is anchored to the plasma membrane at both ends, at least one of the heptad positions *e* or *g* is most probably buried as it is facing the cell surface (Fig. 2C). We therefore reasoned that only heptad positions *b*, *c* and f are more likely to be exposed for interaction with ILT7. Based on these predictions, we selected potentially exposed non-alanine residues, which were part of the quadruple alanine scan null mutants, for individual studies (Fig. 2B-C). Several mutations had a slight to severe ILT7 activation phenotype impairment, but two BST2 mutants, D129A and R136A, completely lost their ability to activate ILT7 (Fig. 3A-B). The defective phenotype of BST2 D129A and R136A mutants was confirmed using the PBMC-based assay. As expected, upon engagement of BST2 WT with ILT7, IFN-I production was significantly reduced in the co-culture (Fig. 3C and S2). In agreement with the lack of ILT7 activation observed using the ILT7 reporter assay, BST2 D129A and R136A mutants were unable to significantly trigger a repression suppression of the IFN-I pathway triggered through TLR7 activation (Fig. 3C and S2). Interestingly, the two mutated residues, D129 and R136 are located in adjacent *c* heptad positions (Fig. 2C) and as such could form a potential ILT7-binding surface in the secondary structure of BST2 (Fig. 3D).

**Fig 2.**
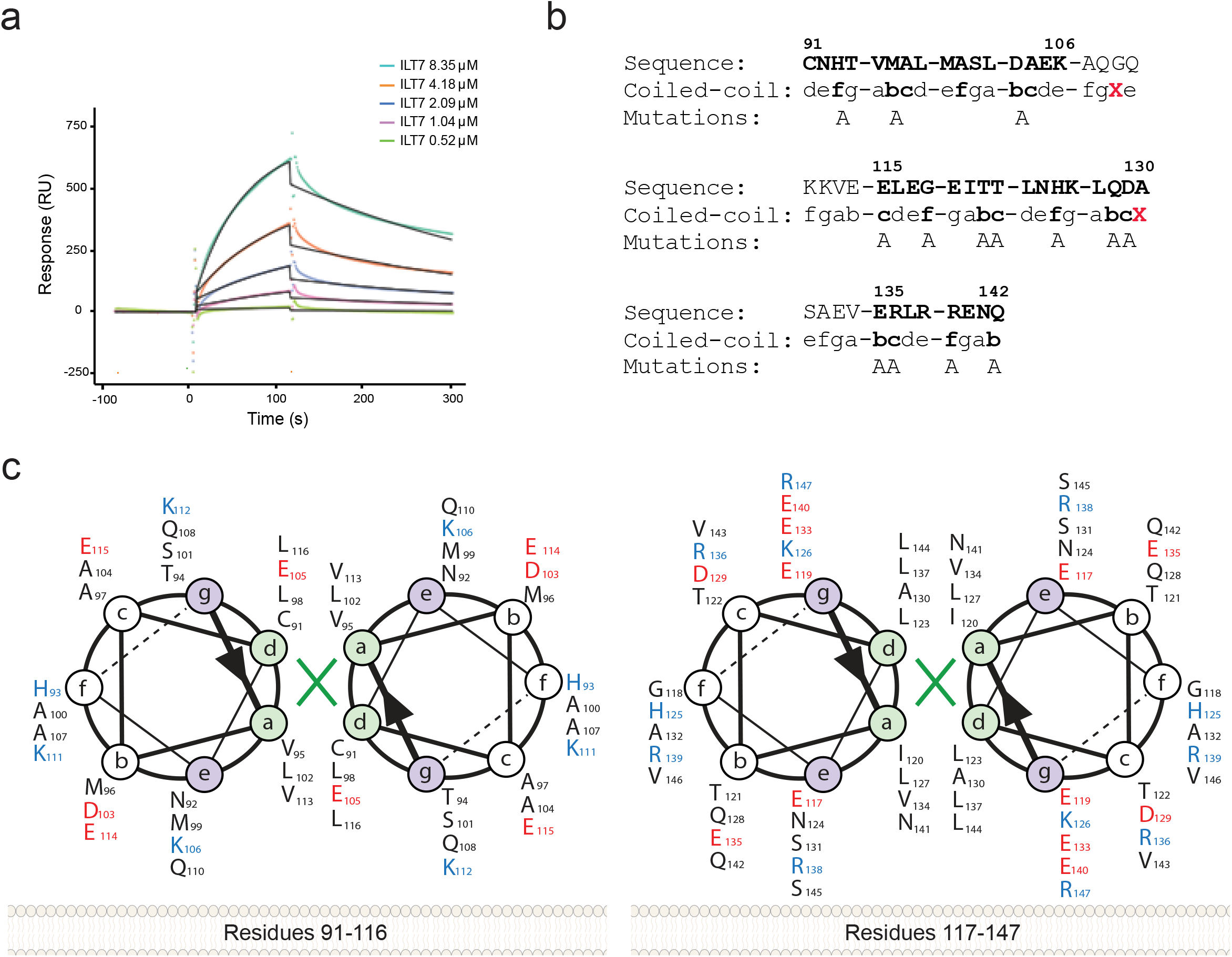
Binding of ILT7 to BST2 coiled-coil region. **a,** Recombinant GST-BST2 (80-147) pre-coated on the surface of Biacore sensor chip, was mixed with the indicated concentrations of bacILT7 24-435. The kinetic response data after subtracting the value from a reference cell coated with GST alone are shown. Kinetic constants (K_D_=2.62 μM, k_on_=0.908×10^3^ M^-1^s^-1^, k_off_ =2.38×10^-3^ s^-1^) were derived by fitting the data (dotted lines) to a 1:1 Langmuir model (black lines) using local R_max_ parameters (chi_2_ =4.95). **b,** Identification of potentially individual exposed residues within BST2 coiled-coil region. Sequence of BST2 coiled-coil region, position of alanine scan mutants that failed to activate ILT7 (in bold), coiled-coil heptad positions (denoted a-g), positions most likely to be exposed (in bold; stutter position in red) as well as individual alanine substitutions of predicted BST2 exposed residues are indicated. **c,** Helical wheel diagram of the homodimeric parallel coiled-coil of BST2 (residues depicted from position 91-116 and 117-147). The primary structure of each helix is characterized by a periodicity of seven residues or heptad repeat pattern. Heptad positions a and d (shaded green) are typically hydrophobic core residues present at the interface of the two helices, while e and g positions (shaded purple) typically form inter-helical ionic interactions and are all involved in the dimer formation. Residues often found in the remaining heptad repeat positions b, c, and f are exposed for potential interaction with binding partners. Charged amino acids are colored (negative in red and positive in blue). Given the BST2 unique double membrane anchored conformation, its fixed membrane facing interphase is highlighted as well.

**Fig 3.**
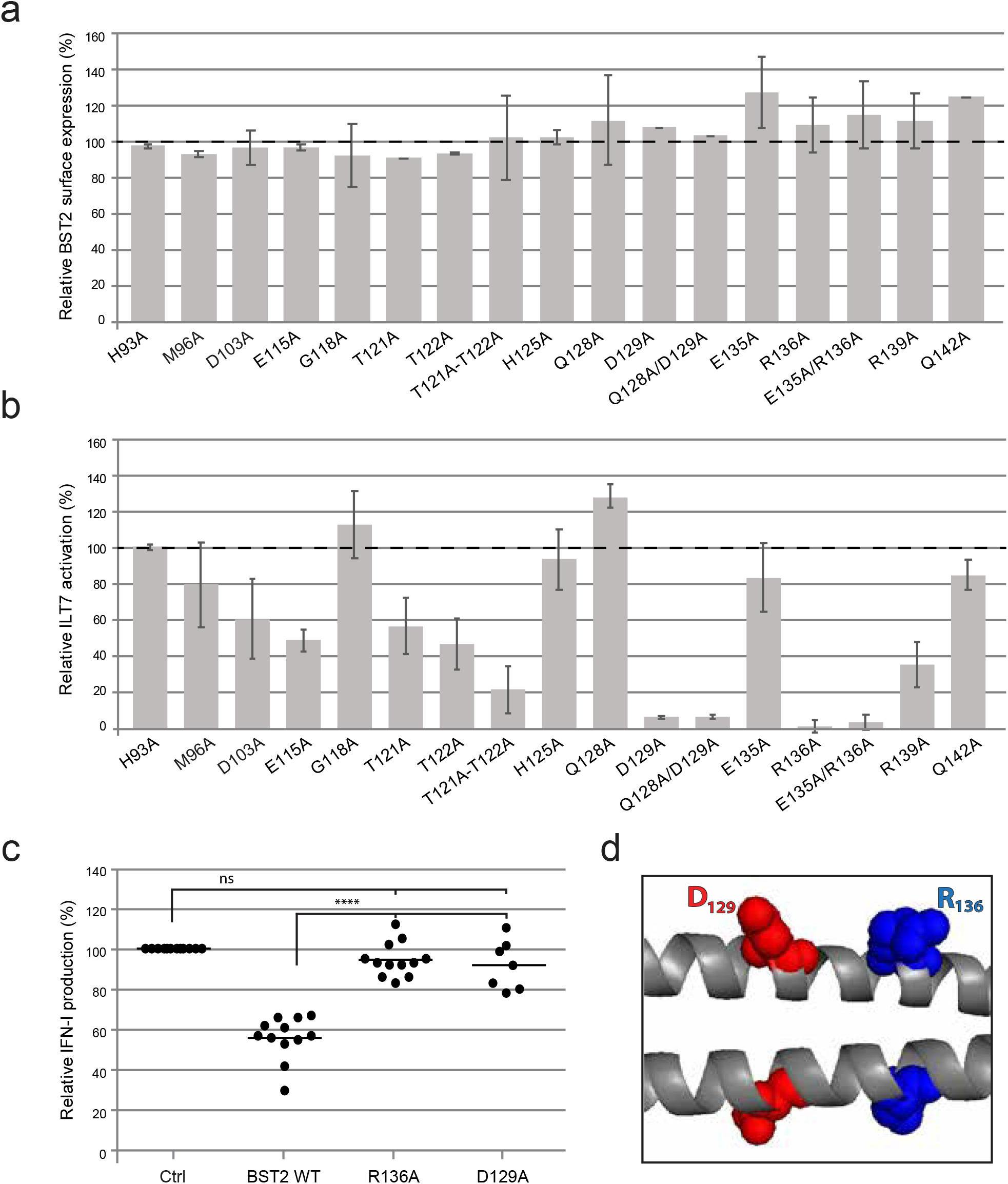
Effect of alanine substitutions of predicted exposed BST2 residues on ILT7 activation. **a,** Relative BST2 surface expression in HEK-293T cells transfected with control plasmids or plasmids encoding the indicated BST2 mutants (n=6). Percentages of MFIs were calculated as in Figure 1. **b,** ILT7+ NFAT-GFP reporter cells were co-cultured for 18-24 h with HEK-293T cells expressing the indicated BST2 mutants and analyzed by flow cytometry (n=6). Percentage of ILT7 activation was plotted as described in Figure 1. Error bars represent SD. **c,** HEK-293T cells expressing the indicated BST2 mutants were co-cultured with freshly isolated PBMCs and levels of bioactive IFN-I released in supernatants in response to TLR7 agonist was measured 18-24 h later. Results are expressed as relative percentage of IFN-I released by PBMCs in contact with HEK-293T cells transfected with the empty plasmid in presence of TLR 7 agonist (100%, n=12). Statistical significance was determined by applying repeated measures ANOVA with Bonferroni’s multiple comparison test.**d,** Putative orientation of residues D129A and R136A extrapolated from the crystal structure of BST2 residues 80–147 published by Hinz, et al.^26^

### BST2 coiled-coil dynamic instability is important to modulate ILT7 activation

Alanine substitutions of residues 63 to 78 in the N-terminal domain of BST2 enhanced ILT7 activation by 3-to 5-fold (Fig. 1D). This same region was previously shown to display pronounced flexibility in the context of BST2 dimers and to act as the core of a putative antiparallel 4-helix bundle made by two BST2 parallel dimers under reducing conditions^24,25^. A single residue mutant in this region, L70D, was sufficient to disrupt tetramers without affecting the formation of BST2 dimers under reducing condition but had only limited effects on BST2 antiviral activity^24^. When tested in the ILT7 reporter assay, substitution of leucine residue at position 70 for aspartic acid significantly enhanced activation (Fig. 4A-B). Interestingly, this mutant behaved just as BST2 WT in the PBMC-based assay (Fig. 4C). Given the structural feature displayed by this region, the L70D mutation could also have long range effects on the dynamic of the coiled-coil region. Crystal structure models from Schubert and colleagues suggest that residue L70 is buried in a hydrophobic core stabilizing the BST2 dimer^24^. It is therefore possible that disrupting a strong hydrophobic core within the N-terminus end of the dimer by adding a charged residue would provide more flexibility to the C-terminus coiled-coil domain. In order to test whether the dynamic instability of the BST2 coiled-coil plays a role in ILT7 activation, we incorporated two cysteines at positions L127 (L127C) and V134 (V134C) predicted to be close enough in the dimer to allow formation of disulfide bonds (Fig. 4D). As predicted, restricting the plasticity of the BST2 coiled-coil region significantly reduced ILT7 activation in the ILT7 reporter assay as well as in the PBMC-based assay (Fig. 4A-C). These results highlight the importance of BST2 coiled-coil characteristic of dynamic instability for ILT7 activation and suggest that the N-terminal domain might impose a structural constraint on the coiled-coil dynamic.

**Fig 4.**
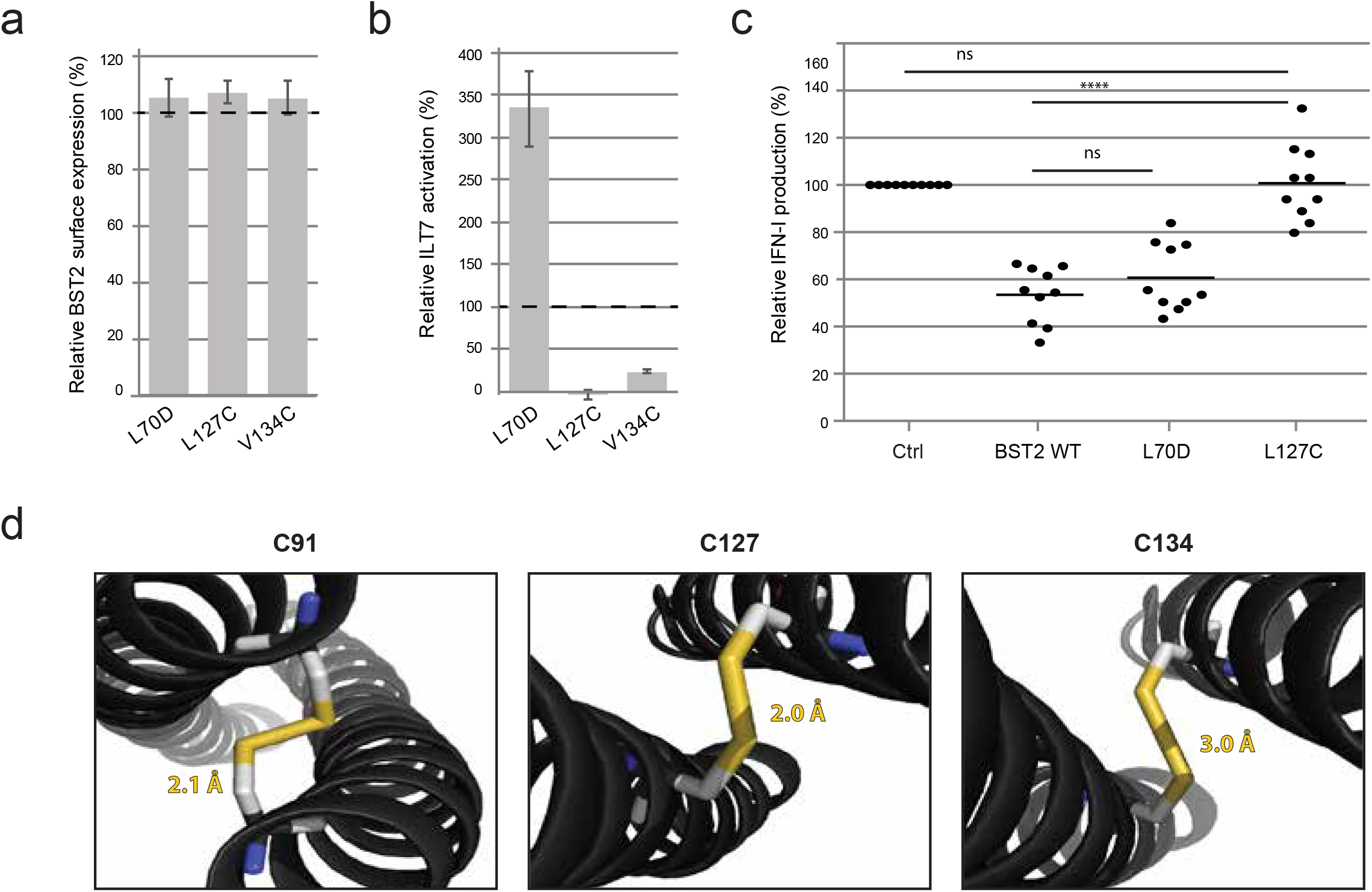
Effect of BST2 L70D mutation and incorporation of engineered di-sulfide bonds on ILT7 activation. **a,** Relative BST2 surface expression in HEK-293T cells transfected with control plasmids or plasmids encoding the indicated BST2 mutants (n=6). Percentages of MFIs were calculated as in Figure 1. **b,** ILT7+ NFAT-GFP reporter cells were co-cultured for 18-24 h with HEK-293T cells expressing the indicated BST2 mutants and analyzed by flow cytometry (n=6). Percentage of ILT7 activation was plotted as described in Figure 1. Error bars represent SD. **c,** Control, BST2 WT or selected mutants (L70D and L127C)-expressing HEK-293T cells were co-cultured with freshly isolated PBMCs and levels of bioactive IFN-I released in supernatants in response to TLR7 agonist was measured as described in Figure 3 (n=10). **d,** Predicted distance between natural cysteine at position 91 on sister BST2 helices as well as between cysteines residues replacing proposed polar dimerization contacts at positions 127 and 134 (in adjacent heptad a positions, see Fig. 2B-C). Cysteine replacement mutations were introduced using Coot and di-sulfide bonds visualized using Pymol. Distances were extrapolated from the crystal structure of BST2 residues 80–147 published by Hinz, et al.^26^ The distance between the sulfurs in the disulfide bond in Å is shown for each of the above-mentioned positions.

### ILT7 activation requires a structurally stable BST2 dimer

Although there are three disulfide bonds (C53, C63 and C91) stabilizing the BST2 dimer^24–26^, none of the three individually is required for BST2 antiviral function^21,22^. Loss of cysteines at position 53 and 63 did not affect the ability of BST2 to activate ILT7 (Fig. 5A-B). On the other hand, loss of cysteine residue at position 91 drastically reduced ILT7 activation. Indeed, any combination that included a mutation at cysteine residue 91 displayed a strong defect in ILT7 activation (Fig. 5A-B). Of note, N-acetylglucosamine residues at position N92 are postulated to be perpendicular to the C91–C91 disulfide bond^24^. Interestingly, while N-glycosylation sites were not required for ILT7 activation, disrupting C91 disulfide bond in absence of glycosylation at position 92, restored BST2-mediated ILT7 activation to BST2 WT levels (Fig. 5A-B). Nevertheless, the stabilizing role of the disulfide bonds was found to be required for ILT7 activation as mutation N92Q could not rescue activation when the three cysteines were mutated (Fig. 5A-B). In agreement with the ILT7 reporter cell assay, BST2 C91A mutant was unable to significantly trigger the ILT7-dependent IFN-I repression pathway in PBMCs, while no significant differences were observed between BST2 WT and the C91A/N92Q mutant (Fig. 5C). These results indicate that although the structural stability of the BST2 dimer conferred by the disulfide bonds is critical for BST2-mediated ILT7 activation, the disulfide bond specifically formed through Cys 91 might also contribute to the proper orientation of a neighboring highly glycosylated site that otherwise could impair BST2-ILT7 engagement.

**Fig 5.**
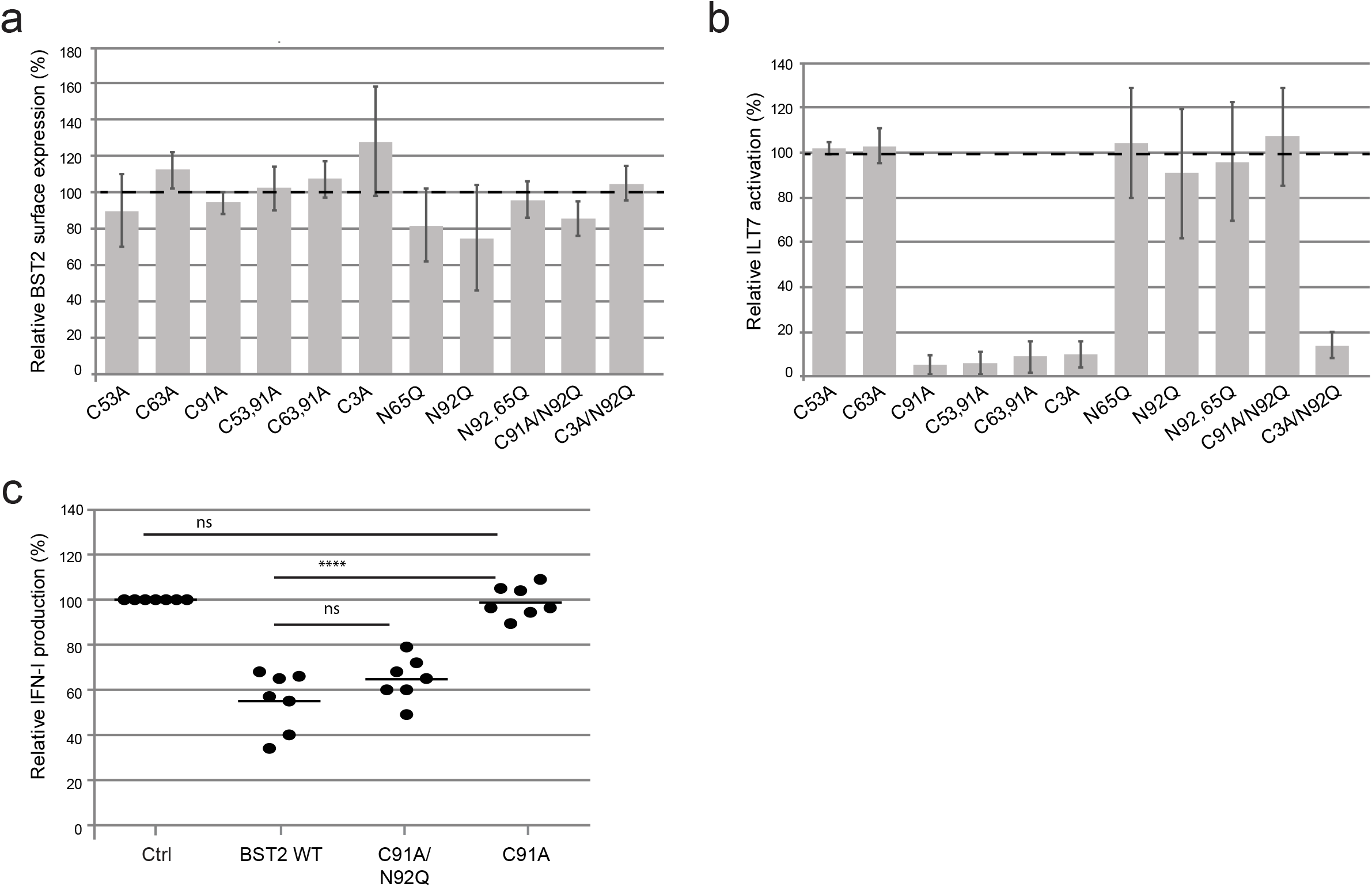
Role of BST2 cysteine or asparagine residues in ILT7 activation. **a,** Relative BST2 surface expression in HEK-293T cells transfected with control plasmids or plasmids encoding the indicated BST2 mutants (n=6). Percentages of MFIs were calculated as in Figure 1. **b,** ILT7+ NFAT-GFP reporter cells were co-cultured for 18-24 h with HEK-293T cells expressing the indicated BST2 mutants and analyzed by flow cytometry (n=6). Percentage of ILT7 activation was plotted as described in Figure 1. Error bars represent SD. **c,** Control, BST2 WT or selected mutants-expressing HEK-293T cells were co-cultured with freshly isolated PBMCs and levels of bioactive IFN-I released in supernatants in response to TLR7 agonist was measured as described in Figure 3 (n=7).

Given that a stable BST2 dimer is required to activate ILT7 and that the ILT7 coreceptor, FcϵRIγ, requires dimerization for proper activation^37^, we investigated whether ILT7 activation requires the engagement of two functionally active units within the BST2 dimer. Using a double inverse gradient of expression of BST2 WT versus the R136A null mutant, we determined that only homodimers of BST2 WT are capable of activating ILT7 while heterodimers between BST2 WT and the R136A mutant are inactive (Fig. 6 A-B, S3A). To support these findings, we demonstrated by co-immunoprecipitation assay that BST2 WT was capable of forming dimers with BST2 mutant R136A as efficiently as with BST2 WT (Fig. S3B). These findings indicate that two functionally active units of BST2 forming a stable dimer are required to activate ILT7.

**Fig 6.**
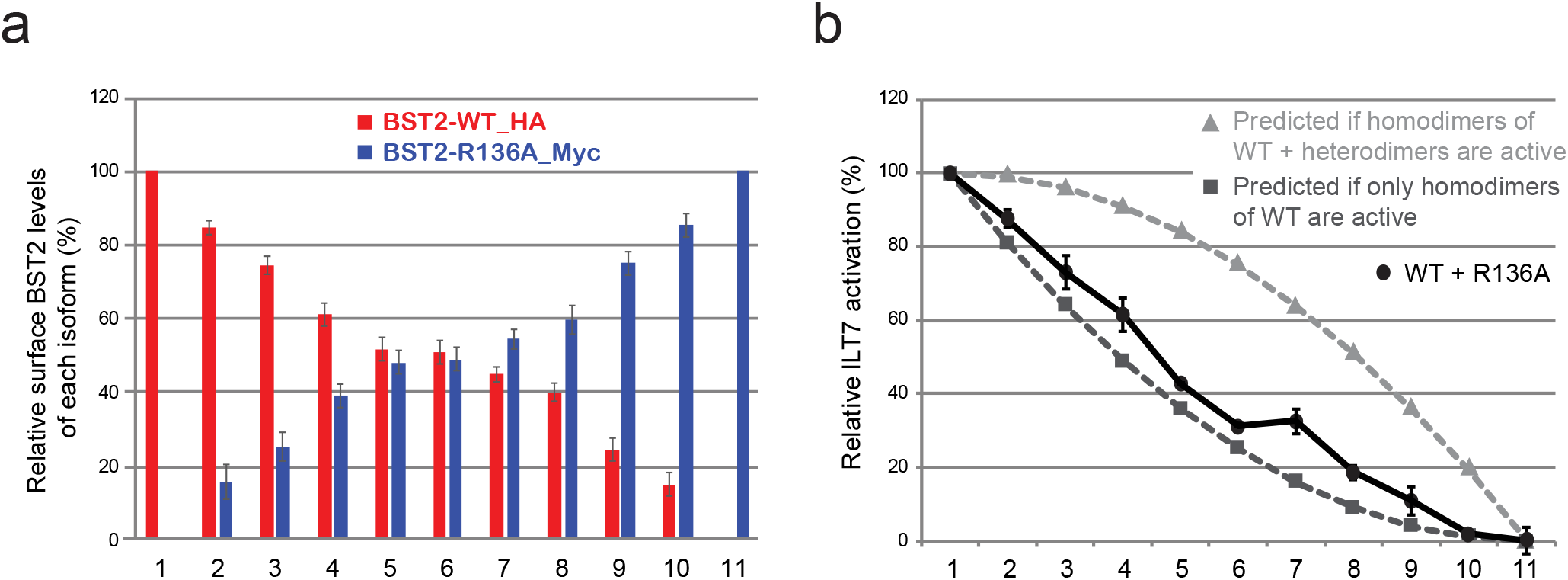
ILT7 activation in response to different ratios of functional and non-functional BST2 isoforms. A converging (or double) gradient of expression of HA-BST2 WT and Myc-BST2 R136A was generated in HEK-293T cells such that conditions 1 and 11 represent solely HA-BST2 WT or Myc-BST2 R136A, respectively whereas condition 6 is the mid-point where each isoform is predicted to be at 50%. Total amounts of transfected DNA were identical in all conditions. **a,** Flow cytometry analysis of surface BST2 in HEK-293T cells transfected with conditions 1 to 11 of the above-mentioned converging gradient for HA-BST2 WT (red) and Myc-BST2 R136A (blue). Ratios of each isoform for all conditions are plotted as percentage of total BST2, as calculated in the examples described in Fig. S3. **b,** Relative ILT7 activation from converging BST2 gradient. ILT7+ NFAT-GFP reporter cells were co-cultured for 18-24 h with control (empty) or HEK-293T cells expressing the above-mentioned BST2 gradient for 18-24 h and analyzed by flow cytometry. Percentage of ILT7 activation was plotted as % of GFP+ cells in each condition relative to the HA-BST2 WT condition (100%) after subtracting the % of GFP+ cells in the no BST2 condition (0%). The predicted curves if only homodimers of BST2 WT or if both homodimers of BST2 WT and heterodimers between BST2 WT and R136A were capable of activating ILT7 are indicated in dark grey squares and light grey triangles respectively. The mean values of four independent experiments are plotted and error bars represent SD.

### Natural variants of BST2 exhibit distinct ILT7 activation phenotypes

To examine if the BST2 functional domains identified so far can be affected *in vivo*, several natural variants of BST2 were analyzed, including single nucleotide polymorphisms (SNP) and somatic mutations associated to specific cancers (Fig. 7A, Table S1). Most of the SNP tested were comparable to BST2 WT for ILT7 activation; however, three SNPs, namely D103N, E117A and D129E, the latter revealing a substitution at the critical D129 residue for a conserved charged amino acid, exhibited a partially impaired ILT7 activation phenotype (Fig. 7B). Likewise, three of the somatic mutations tested (E62G-hepatocellular carcinoma, Q110H-ovarian cancer and S145R-lung cancer) activated ILT7 to the same extent as BST2 WT. Interestingly, one somatic mutation (E119K-melanoma) located in the coiled-coil domain was completely defective while three somatic mutations identified in distinct cancer tissues (C63Y-uterine cancer, Q87H-colon cancer and T90P-breast cancer) and mapping outside of the coiled-coil structure exhibited enhanced activation capacities (Fig. 7B). In agreement with the ILT7 reporter assay, mutation E119K was completed defective in the PBMC assay (Fig. 7C). Based on its heptad *g* position (Fig. 2C), it is predicted that the charge alteration at this position would have a very detrimental effect on the overall coiled-coil stability, further highlighting the importance of the integrity of this structure for ILT7 activation. However, in contrast to the BST2 C63Y and T90P mutants that behaved just as WT in this assay, BST2 Q87H strongly repressed IFN-I production (Fig. 7B-C). Taken together these results suggest that variants of BST2 with opposed potential to activate ILT7 can be found *in vivo* and be selected under pathological malignant conditions.

**Fig 7.**
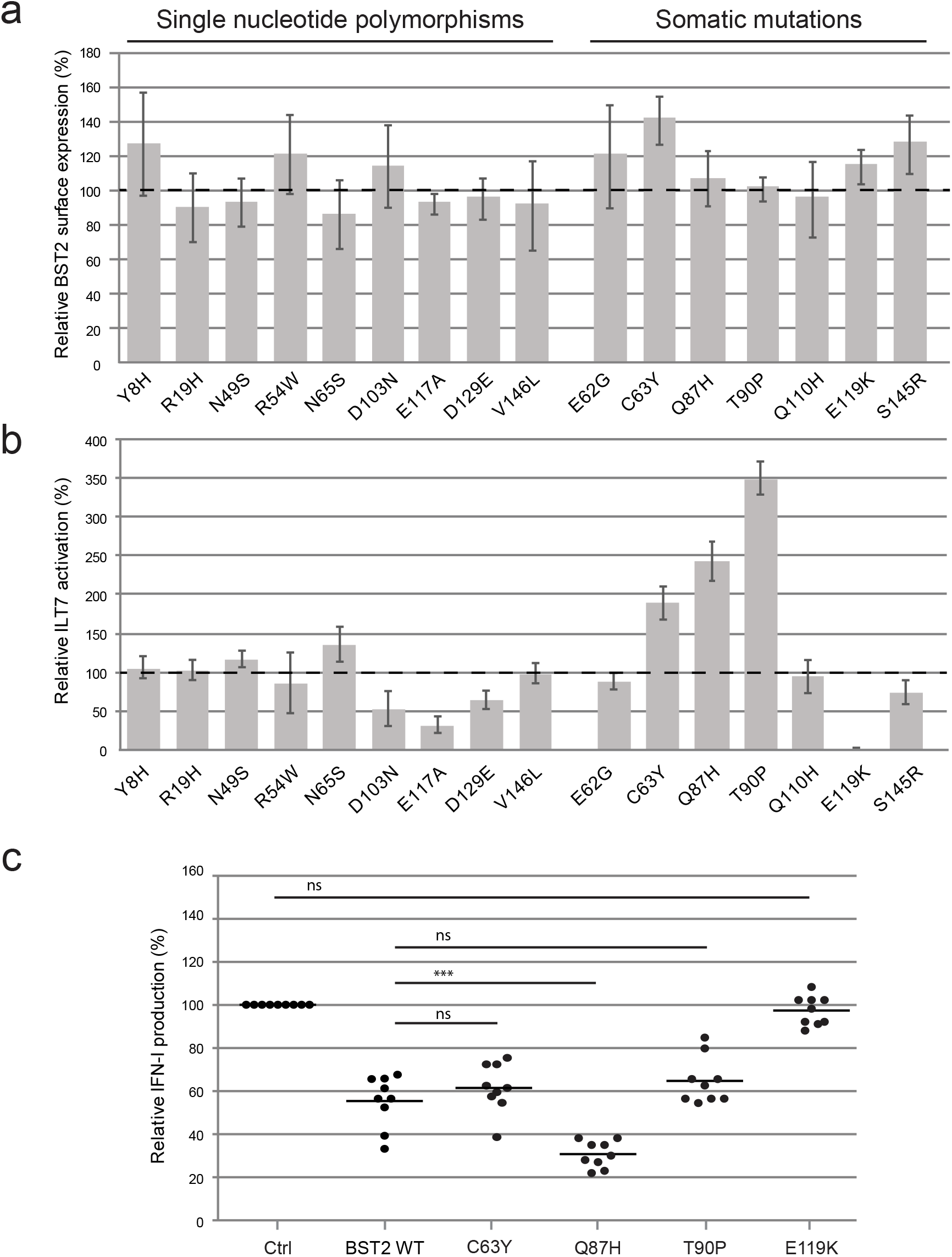
Effect of natural variants of BST2 on ILT7 activation. **a,** Relative BST2 surface expression in HEK-293T cells transfected with control plasmids or plasmids encoding the indicated BST2 mutants containing the single nucleotide polymorphisms or somatic mutations described in table S1 (n=6). All Percentages of MFIs were calculated as in Figure 1. **b,** ILT7+ NFAT-GFP reporter cells were co-cultured for 18-24 h with HEK-293T cells expressing the indicated BST2 mutants and analyzed by flow cytometry (n=6). Percentage of ILT7 activation was plotted as described in Figure 1. Error bars represent SD. **c,** Control, BST2 WT or selected mutants-expressing HEK-293T cells were co-cultured with freshly isolated PBMCs and levels of bioactive IFN-I released in supernatants in response to TLR7 agonist was measured as described in Figure 3 (n=9). Statistical significance was determined by applying repeated measures ANOVA with Bonferroni’s multiple comparison test was used.

### Post-binding events associated to BST2 coiled-coil plasticity are required to activate ILT7

To further understand the molecular mechanism underlying ILT7 activation by BST2, we analyzed the binding strength of selected BST2 mutants to ILT7 using microscale thermophoresis (MST). The recorded binding strength of BST2 WT ectodomain to ILT7 ectodomain by MST (Fig. 8 and Fig. S4, K_D_=2.45 ± 0.3 μM) was remarkably similar to that we previously reported using surface plasmon resonance (SPR) (K_D_=2.33 μM^30^). Mutations in the postulated ILT7-binding surface, BST2 D129A and R136A, that failed to activate ILT7 in the reporter assay and in the PBMC-based assay exhibited no binding to ILT7 by MST, validating these residues as critical for ILT7 binding (Fig. 8A and Fig. S4A). Interestingly, mutation L127C, which prevented BST2 activation of ILT7 in both functional assays by presumably limiting the plasticity of the coiled-coil, interacted with ILT7 with comparable strength than BST2 WT (Fig. 8B and Fig. S4B, K_D_ =6.45 ± 1.2 μM), implying that binding to ILT7 is required but not sufficient to induce activation of the inhibitory receptor. MST analysis of BST2 mutants that enhanced ILT7 activation in the reporter assay (L70D, T90P and Q87H) and in the PBMC assay (Q87H) revealed that none of these mutants displayed enhanced binding affinities. Indeed, these three mutants had K_D_ values in the lower micromolar range as BST2 WT, although with different thermophoresis (Fig. 8C-D and Fig. S4C-D, Q87H K_D_=5.06 ± 1.1 μM; L70D K_D_ =5.55 ± 1 μM; T90P K_D_=4.49 ± 0.9 μM). Whereas BST2 WT and the BST2 Q87H mutant displayed a difference in amplitude when in complex with ILT7, BST2 mutants L70D and T90P showed a difference in signal direction. Chemical cross-linking of constructs comprising the recombinant ectodomain revealed that all mutants tested still dimerized as indicated by the appearance of new bands migrating at molecular weights between 25 and 35 kDa (Fig. S4E), as previously reported^26^. To analyze the impact of the mutations on the folding and stability of the proteins, we examined the circular dichroism (CD) profiles of selected BST2 mutants and determined their thermostability (melting temperature, Tm). CD analyses revealed that all mutants are highly α-helical in solution, with a propensity to form alpha helices either very similar to BST2 WT (Q87H, L127C) or just slightly reduced (70-90% helical content remaining for D129A, R136A, L70D and T90P; Fig. S5, left panels). Analysis of the thermostability showed that while most mutants (D129A, L70D, Q87H and T90P) displayed melting temperature similar to BST2 WT (~61 °C), mutants harboring substitutions at residues L127 or R136 in the coiled-coil domain exhibited a reduced thermostability with Tm values of ~53 °C and ~54 °C, respectively (Fig. S5, right panels). Despite the lower helical content and reduced thermostability of some of the mutants, none of these mutations completely lost the integrity of the coiled-coil structure, as observed for instance under reducing condition when the Tm drops to as low as ~35 °C ^26^. Taken together, these results validate that post-binding events are governing the extent of ILT7 activation by BST2.

**Fig 8.**
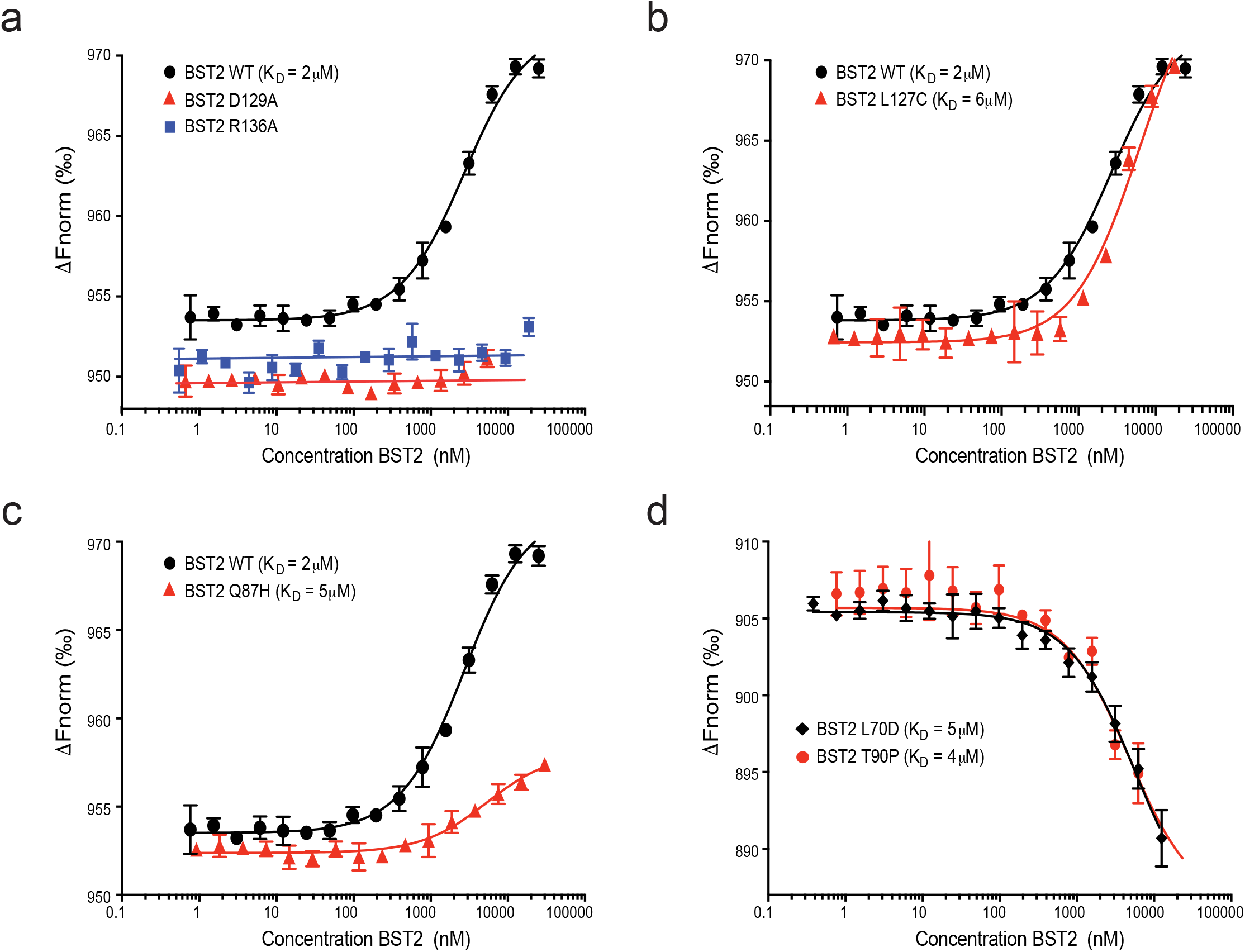
Thermophoretic analysis of BST2–ILT7 interaction. A series of 16 dilutions of recombinant BST2 (residues 47-159) WT or mutants were mixed with 1µM of bacILT7 24-223. After 2min incubation the samples were loaded into Monolith NT.115 Capillaries. Dose-response curve of ILT7 towards **a,** BST2 WT or mutants D129A and R136A, **b,** BST2 WT or mutant L127C, **c,** BST2 WT or mutant Q87H, and **d,** mutants L70D and T90P were generated. All experiments were done in triplicates. All resulting dose-response curves were fitted to a one-site binding model to obtain K_D_ values (indicated in graphs). Error bars indicate SD. MST experiments were performed at a LED power of 80% and at medium MST power. Fnorm = normalized fluorescence.

## Discussion

The interaction of the human pDC receptor ILT7 with its ligand, BST2, has important biological implications as it significantly regulates pDC’s TLR-induced innate immune responses during inflammatory injuries, a function that can be diverted in disease states^11,30,38^. However, the structural determinants and functional features of BST2 that govern ILT7 engagement and activation are still largely undefined. Our mutagenic studies of the BST2 ectodomain combined with biophysical analysis of recombinant mutant molecules identify two structurally distinct regions of the BST2 ectodomain that play divergent role during activation of ILT7. While the coiled-coil region of BST2 is found to contain amino acid residues required for binding ILT7, the flexible N-terminal region of the ectodomain appears to act as a negative modulator of BST2-mediated ILT7 activation. Through analysis of specific mutants, we further demonstrate that binding of BST2 to ILT7 is not sufficient for activation of the receptor and provide evidence that post-binding structural rearrangements involving the coiled-coil structure are likely required to optimally trigger ILT7 signaling. Importantly, our results highlight the biological relevance of these newly defined domain of BST2 by establishing that mutations with opposed potential to activate ILT7 can be found *in vivo* and be selected under pathological malignant conditions.

The human BST2 coiled-coil is characterized by an inherent dynamic instability, which has been proposed to provide structural plasticity during dynamic events such as viral assembly and release^24–27^. Indeed, the integrity of the coiled-coil structure rather than its amino-acid sequence composition was found critical for optimal BST2 antiviral activity^22,32,33^. However, the presence of large patches of amino acid sequence conservation between human BST2 and its counterparts encoded by other primates suggest that the structure could also be important for a conserved cellular function involving a cell surface binding partner^27^. A role of the BST2 coiled-coil as a binding interface for the ILT7 receptor is indeed supported by our *in vitro* binding studies revealing that this structure is sufficient for binding ILT7 as well as by our alanine scan mutagenesis, which clearly demonstrate that mutations encompassing the coiled-coil drastically affect ILT7 activation. Targeted mutagenesis of amino-acid residues predicted to be exposed within the coiled-coil heptad repeats identify well conserved residues D129 and R136 as essential on their own for binding ILT7 and as expected inducing its activation. In the case of the aspartic acid residue at position 129, substitution for a similarly charged glutamic acid, was found still detrimental for ILT7 activation, pointing to a critical role of an aspartic residue at that position. The fact that the overall integrity of the coiled-coil appears minimally affected in both mutant, as shown by our biophysical analyses, and that they retain the ability to form dimers, suggests that these exposed residues in adjacent c positions might represent an ILT7-binding surface in BST2 although we cannot exclude the possibility that ILT7 contacts BST2 at two distinct sites. Interestingly, the requirement of two, rather than one, binding contacts was previously reported for ILT2 (also known as LIR-1 or LILRB1), another member of the ILT family, which binds HLA-A2^39^.

Previous structural and functional studies have highlighted the critical role of disulfide linkages in stabilizing the BST2 dimer as well as the necessity for disulfide cross-linking through at least one of the three conserved cysteines for BST2 antiviral activity^21,22,24–27^. Our results demonstrate that the stabilizing role of disulfide bonds is required for ILT7 activation. They also reveal an essential role of cysteine 91 as disulfide cross-linking at this position is predicted to prevent the detrimental effect of a neighboring asparagine residue on the BST2-mediated ILT7 activation. Interestingly, none of the other two cysteines, C53 and C63, were found to individually impact ILT7 activation, indicating that disulfide cross-linking at C91 might serve two functions: first, ensuring the proper structural orientation of the C-terminus of the ectodomain, and, second stabilizing the labile coiled-coil during dynamic processes of disassembly and reassembly, which confer a degree of conformational flexibility/plasticity to the molecule^23^. Restricting the dynamic process of the BST2 coiled-coil with engineered disulfide bonds predicted to be formed at positions 127 and 134 drastically impaired ILT7 activation without however impairing binding of BST2 to the receptor (at least with L127C). These results suggest that while binding of BST2 to ILT7 is required for induction of ILT7 signaling, it might not be sufficient. This phenotype raises the possibility that post-binding events associated to the plasticity of the BST2 coiled-coil might be required for ILT7 activation, although we cannot entirely exclude that the lack of ILT7 activation might result from the slightly reduced thermostability of the L127C mutant.

Alanine scan and targeted mutagenesis of the N-terminal domain of BST2 at amino acid residues 63, 70 87 and 90 leads to an enhancement of ILT7 activation in the reporter assay without any major changes in the strength of binding to ILT7 (at least for the L70D, Q87H and T90P mutants). These results suggest that this region of BST2 might exert a structural constraint on the ectodomain of the molecule. They also support the notion that besides binding, other molecular events might qualitatively impact BST2-mediated ILT7 activation. Structural studies of BST2 reveal that the N-terminal domain of BST2 forms a continuous helix with the coiled-coil with and inherent constrained flexibility at a connecting hinge region, which is revealed as a bent structure by small angle X-ray scattering analysis^26,27^. However, the presence of both membrane anchors might constrain the ectodomain into one defined conformation. Thus, disrupting a strong hydrophobic core close to the N-terminus end of the dimer (L70D mutation) could provide more flexibility to the C-terminus coiled-coil portion by reducing the backbone torsion downstream. Similarly, introducing a sharp helix break with a proline before the start of the coiled-coil region (T90P mutation), physically disconnecting the two domains, might relieve this structural constrain and favor ILT7 activation.

On the basis of our analysis of the structure-function relationship of BST2 and the current literature, we propose the following model of BST2-mediated activation of the ILT7 pDC receptor. Two functionally active units of BST2 forming a stable dimer are required to activate ILT7. Upon binding of each BST2 molecules to a single monomeric ILT7/FcϵRIγ pair at either side of the dimeric coiled-coil, the two ILT7/FcϵRIγ pairs are brought close enough to induce their dimerization, a process that is facilitated by the conformational flexibility of the coiled-coil structure. Once close enough, a potential disulfide bond formation linking the two FcϵRIγ co-receptors would stabilize the entire complex as observed for the Fc receptor complex^37^. In this context, mutations D129A or R136A would prevent the initial binding while mutation L127C would limit the conformational flexibility of the coiled-coil required to displace the ILT7/FcϵRIγ complexes close enough for dimerization and subsequent signaling to occur. Relieving the structural constrain exerted by the N-terminal on the ectodomain, as exemplified by mutant L70D or T90P, would make this step more efficient by lowering the energy transition required to mediate this conformational rearrangement, thus allowing a more effective activation of the ILT7 receptor. While this phenotype is observed in the reconstituted ILT7+ NFAT-GFP reporter cell assay (indicator cells are in are mouse hybridoma cell background), which might be more sensitive to structural changes that lower energy transitions, it was not detectable in the more physiological PBMC assay where formation of an active dimeric ILT7/FcϵRIγ complex is likely more efficient and stable. Interestingly, a somatic mutation where glutamine 87 in the flexible connecting region was substituted for a histidine induced a significant increase of the ILT7 activation potential of BST2 in both assays, making it unlikely that this mutation caused this phenotype by lowering the energy transition of the predicted rearrangement step. The fact that the Q87H mutant binds ILT7 with an affinity comparable to WT BST2, rather suggests a potential role of the flexible connecting domain in modulating the quality of the BST2/ILT7 binding. Since the binding affinity of BST2 for the ILT7 receptor is rather low (K_D_ ~2.5 µM), clustering of BST2 of a higher order might be required to fully trigger the ILT7 signaling cascade. Small-angle X-ray scattering (SAXS) analysis of human and mouse BST2 revealed features indicative that the BST2 ectodomain can form higher order associations in solution^27^. Accumulation of BST2 within plasma membrane microdomain at the point of contact with pDC would support the formation of such high order associations and mutations facilitating or limiting the magnitude of this clustering might impact BST2-mediated ILT7 activation. In support of this, cryo-EM studies demonstrate the clustering of BST2 at HIV-1 budding sites in the absence of viral antagonist^40,41^. Yet, despite HIV-1 counteractions, which result in a reduction of surface BST2 and efficient viral particle release, BST2 is still incorporated into virions.^22,42^ These observations provide indirect evidence that conditions not favoring the formation of high order associations render BST2 ineffective at restricting HIV-1 release.

The biological importance of ILT7 activation by BST2 has been difficult to study *in vivo* since this function is not conserved between human and mice, as there is no known murine ortholog for ILT7^14^. Moreover, although mouse and human BST2 ectodomains have an evolutionarily conserved unstable coiled-coil design, there is no actual amino acid sequence conservation^27^ and the newly identified putative ILT7-binding surface is absent in rodent BST2. Indeed, paradoxically, BST2 knock-out mice showed reduced IFN-I secretion by murine pDCs^43^. Several previously identified naturally-occurring SNPs of BST2 in the human populations were investigated in the context of BST2 antiviral restriction and signaling. None of these sequence variants affected the ability of BST2 to restrict HIV-1 release while one variant, Q19H, selectively abolished the NF-κB-dependent signaling activity of BST2^44^. Most of these very rare missense variants of BST2 activated ILT7 as well as BST2 WT, thus highlighting the high conservation of this function in healthy individuals. On the other hand, it is well established that elevated BST2 levels are observed in many types of malignant cells^15^. The elevated BST2 mRNA levels observed in some metastatic and invasive tumors are a strong predictor of tumor size, aggressiveness and often associated with poor patient survival^45,46^. Nonetheless, BST2 is not elevated in all cancer tissues as significant downregulation was noted in particular cases, including B-cell acute lymphoblastic leukemia, liver, and prostate cancer^15,47^. Functional analyses of a selected number of BST2 ectodomain variants enriched in malignant cells when compared to healthy cells from the same individual reveal that most somatic mutations had no impact on BST2 activation except for two sequence variants, E119K and Q87H. The E119K variant found in melanoma abrogates ILT7 activation most likely by affecting the integrity of the coiled-coil structure. In contrast, the Q87H variant associated with colorectal cancer enhances the ILT7 activating capacity of BST2. Thus, these variants could confer evolutionary advantages to the malignant tumors they are associated with by enabling them to modulate pDC’s IFN-I responses and hence either escape IFN-I anti-tumor activity (Q87H) or promote IFN-I pro-tumorigenic potential (E119K)^8,48^.

The IFN system can mount an early and extremely powerful antiviral and antitumor response. By modulating the immune response at its foundation, IFNs can widely reshape immunity to control chronic infectious diseases and invading malignancies. A better understanding of the intimate interplay between BST2 and ILT7 can open new avenues for targeted drug design in the context of anti-viral or anti-cancer strategies.

## Material and Methods

### Antibodies and reagents

Rabbit polyclonal anti-BST2 serum was previously described (Dube, 2010). Mouse monoclonal antibodies (mAb) anti-HA (HA.11 Clone 16B12), and anti-ILT7_alexa647 were acquired from Biolegend. Rabbit anti-Myc Abs and Protein A-HRP were purchased from Sigma and Southern Biotech, respectively. All secondary Abs used for flow cytometry were purchased from Life Technologies. TLR7 agonists Gardiquimod (final concentration: 2.5 μg/ml) was obtained from InvivoGen.

### Cell lines and plasmids

HEK-293T and HEK-blue human IFN reporter cell lines were obtained from ATCC and InvivoGen, respectively. These cells were maintained in DMEM supplemented with 10% FBS. The ILT7 NFAT-GFP (CT550) cells were described previously^10^. These cells were maintained in RPMI supplemented with 10% FBS. High Five insect cells were maintained at a cell density of 0.5–1 × 10^6^ cells/mL in Express Five medium (Life Technologies) supplemented with 16mM L-glutamine. HEK-293T cells were transiently transfected using lipofectamine 2000 (Invitrogen). Usually, 0.5-1.5 μg of DNA was added to each well of a 24-well plate (wp) (125,000 cells/well) and media was replaced 18-24 h post transfection. Unless specified otherwise, un-tagged BST2 open reading frames were cloned into the pcDNA3.1 backbone for transient transfections. Tagged versions of BST2 WT ORF (with internal tags added after amino acid 154) plus cloning restriction enzymes sites were chemically synthesized (Invitrogen) and cloned in the pcDNA3.1 backbone. All mutations introduced in BST2 were generated by PCR-based mutagenesis using specific primers (Table S2). All mutations were validated by automated sequencing. BST2 alanine scanning mutants (cloned in pFLAG-tetherin, with N-terminal tag) were a kind gift from Dr. Paul Spearman^32^. Each mutant in the panel incorporates four alanine residues, starting from amino acid position 47 and continuing through to position 150.

### Surface antigen staining and flow cytometry analysis

BST2 cell-surface staining and flow cytometry analysis was performed as previously described^49^. Briefly, HEK-293T cells were washed in PBS and stained with the specific primary antibody (anti-BST2 rabbit serum, anti-HA or anti-Myc) for 45 min at 4°C. Cells were then washed and stained using appropriate Alexa Fluor-coupled secondary Abs for 30 min at 4°C. After an additional wash, cells were analyzed for cell-surface BST2 expression by flow cytometry. Cells from co-cultures between transfected HEK-293T and ILT7+ NFAT-GFP reporter cells were collected, washed in PBS and stained with anti-ILT7_alexa647 antibodies for 45 min at 4°C. Cells were then washed and analyzed for cell-surface ILT7 and GFP expression by flow cytometry. Fluorescence intensities were acquired using a Cyan ADP flow cytometer and data was analyzed using the FlowJo software (Treestar). In all histograms shown, mean fluorescence intensity (MFI) values are shown for each sample.

### Activation of ILT7 using ILT7^+^ NFAT-GFP reporter cells

Two days prior to co-culture, HEK-293T cells were transfected with empty pcDNA3.1, and plasmids encoding for BST2 WT or mutants. ILT7^+^ NFAT-GFP reporter cells (100,000 cells/well) were added in a final volume of 500μl. The co-cultures were maintained for an additional 18-24 h, at which time samples were analyzed by flow cytometry for surface ILT7 and % of GFP expressing reporter cells from the ILT7^+^ gate. Given that the extent of ILT7 activation is directly proportional to surface levels of BST2 on co-cultured 293T targets cells^30^, expression of all BST2 mutants was standardized to match that of the internal BST2 WT control.

### Preparation of PBMCs and co-cultures

Peripheral blood samples were obtained from healthy adult donors who gave written informed consent in accordance with the Declaration of Helsinki under research protocols approved by the research ethics review board of the IRCM. PBMCs were isolated by Ficoll-Paque centrifugation (GE Healthcare) and cultured in RPMI-1640 media supplemented with 10% FBS. Two days prior to co-culture, HEK-293T cells were transfected with empty pcDNA3.1 or plasmids encoding for BST2 WT or mutants. PBMCs at a ratio of 3:1 (PBMC:293T cell) were added in a final volume of 250μl. After 4 h of co-culture, TLR7 agonist Gardiquimod was added to a final concentration of 2.5 μg/ml and cells were kept in co-culture for an additional 18-24 h as previously described^30^. In all conditions, co-cultures were then transferred to a V-bottom 96-well plate and centrifuged for 5 min at 400g. As a control, transfected HEK-293T cells were treated with TLR7 agonist in absence of PBMCs. Supernatants were then used to quantify the amounts of bioactive IFN-I produced. Each experimental replicate (n) was performed using cells from a different donor.

### Quantification of IFN-I

Detection of bioactive human IFN-I was performed using reporter cell line HEK-Blue IFN-α/β (InvivoGen) as previously described^12^. IFN-I concentration (U/ml) was extrapolated from the linear range of a standard curve generated using known amounts of IFN-I.

### Surface plasma resonance

For Surface Plasmon Resonance (SPR) analysis, the synthetic codon-optimized ILT7 gene (Eurofins Genomics) encoding residues 24 to 435 (bacILT7 24-435) was cloned into the transfer vector pFL, followed by Tn7-based transposition into the EMBacY bacmid to generate a recombinant baculovirus^50^. The extended coiled-coil domain of BST2 (residues 80 to 147) was cloned into the pBADM30 expression vector in order to construct a His-tagged GST-fusion protein. The supernatant of SF21 insect cells secreting bacILT7 was collected 4 days post infection, dialyzed against buffer A (20 mM Tris, 150 mM NaCl, pH 7.2) and concentrated 2-fold. The supernatant was then applied to a nickel-affinity chromatography (Qiagen) column. The column was washed sequentially with buffer A containing 10, 50 and 70 mM imidazole, followed by elution of bacILT7 with buffer A containing 300 mM imidazole. The eluted fractions were pooled and dialyzed extensively against buffer B (20 mM Tris, 150 mM NaCl, 10% glycerol). Analytical size exclusion chromatography showed that the majority of the protein eluted in a peak at 13 mL from a Superdex 200 column (GE Healthcare). BacILT7 was dialysed against HBS-PE and cleared by centrifugation at 100,000 g for 20 min. GST-BST2 (80-147) was expressed in E. coli Rosetta (DE3) cells (Novagen) and purified in buffer C (20 mM Tris, 100 mM NaCl, pH 7.5) by nickel-affinity chromatography (Qiagen) followed by size-exclusion chromatography on a Superdex 200 column in buffer D (20 mM HEPES, 100 mM NaCl, 10 mM EDTA, pH 7.5). SPR was performed on a Biacore 3000 (GE Healthcare) system using HBS-PE as a running buffer (10 mM HEPES pH 7.5, 150 mM NaCl, 3 mM EDTA and 0.005% Tween-20). GST-BST2 (80-147) was diluted to 5 µg/ml in 10 mM sodium acetate (pH 4) buffer and covalently immobilized to the surface of a CM5 sensor chip by amine coupling according to the manufacturer’s instructions, yielding an Rligand of 6550 RU. A reference flow cell was generated by amine coupling of GST alone (R_ligand_=1028). BacILT7 was serially diluted into running buffer and passed over the chip at a flow rate of 10 µl/min. The response from the GST-coated reference cell was subtracted from the response resulting from specific binding to the target protein. Regeneration of the sensor chip was achieved with 10 mM HCl for 60 seconds. The spikes present in the sensorgrams are due to a delay in the bulk refractive index change between the flow cells, which is exacerbated by the 10 µl/min flow rate. Data were analyzed with the BIAevalution software version 4.1.

### BST2 dimerization assay

A converging (or double) gradient of expression was generated by transfecting different ratios of plasmids encoding HA-BST2 WT and Myc BST2 136A mutants such that condition 1 and 11 represent only HA-BST2 WT or Myc-BST2 R136A, respectively. and condition 6 is the mid-point where each plasmid was transfected at equal ratios (50% of each). For each condition, 1.2 μg of total DNA was transfected. The ratio of each expression plasmid varied by 10% between each condition such that as the HA-BST2 WT-expressing plasmid was reduced, the Myc-BST2 R136A-expressing plasmid was increased. Forty-eight hours post transfection, a replica well was used for co-culture with ILT7^+^ NFAT-GFP reporter cells to measure ILT7 activation as described above. In parallel, another replica well was used for surface BST2 staining using either anti-HA, anti-Myc or anti-BST2 antibodies followed by flow cytometry analysis. Staining for total BST2 was used as an internal control to ensure that similar levels of total surface BST2 were achieved in for all transfection conditions

### Co-immnuoprecipitation and Western blot

HA-CAML-or HA-BST2 WT-expressing plasmids were cotransfected with Myc-tagged BST2 (WT or R136A mutant) expressors in HEK-293T. Cells were harvested and lysed in RIPA-DOC buffer (10 mM Tris pH 7.2, 140 mM NaCl, 8 mM Na_2_HPO_4_, 2 mM NaH_2_PO_4_, 1% Nonidet-P40, 0.5% sodium dodecyl sulfate, 1.2 mM deoxycholate) 48 h post-transfection. Ten percent of each lysates was preserved to control for protein expression (input). The remaining cell lysates were incubated with anti-HA for 2 h at 4°C, prior to precipitation with Protein A Sepharose beads (GE Healthcare). Immunoprecipitates were separated by 12.5% SDS-PAGE and analyzed for the presence HA-BST2 WT or Myc-BST2 (WT or R136A mutant) or the negative control HA-CAML by western blot using either anti-HA or anti-Myc Abs.

### Microscale thermophoresis

ILT7 24-223 was expressed and purified as indicated above. ILT7 eluting in the central fraction from the gel filtration column corresponding to monomeric ILT7 was used for all microscale thermophoresis experiments at 1.1 µM in a buffer containing 20 mM HEPES pH7,5, 150mM NaCl. The BST2 ectodomain (residues 47 to 159), WT or mutants, were cloned into expression vector pPROEX HTb (Invitrogen). BST2 protein expression was performed in E. coli Rosetta2 cells induced with IPTG at 20°C overnight. Cells were lysed in buffer A (20mM Tris pH 8.0, 100mM NaCl, 10mM imidazole). Proteins were purified by nickel-affinity chromatography, and washed with buffer B (20mM Tris pH 8.0, 100mM NaCl) containing 20mM and 50mM imidazole for the first and second wash. Only one wash was performed for the BST2 L70D mutant purification. The bound protein fraction was eluted with buffer B containing 250mM imidazole. The His-tag was removed by TEV protease cleavage and both TEV and uncleaved protein were removed by nickel-affinity chromatography. Final purification steps included a size-exclusion chromatography (Superdex 75, GE Healthcare) in buffer C (20 mM HEPES pH 7,5, 150 mM NaCl). The supernatant of High Five insect cells secreting bacILT7 was collected 4 days post infection, dialyzed against buffer A (20 mM Tris, 150 mM NaCl, pH 7.2). The supernatant was then applied to a nickel-affinity chromatography (His Trap excel, GE Healthcare) equilibrated with buffer B (20mM Tris pH 7.2, 150mM NaCl) and washed with buffer B containing 30mM imidazole. The bound protein fraction was eluted with buffer B containing 500mM imidazole and 10% glycerol. Final purification step included a size exclusion chromatography (Superdex 200 increase column, GE Healthcare) in buffer C (20mM HEPES pH7,5, 150mM NaCl). ILT7 purified protein was labeled using the Protein Labeling Kit RED-NHS (NanoTemper Technologies). The labeling reaction was performed according to the manufacturer’s instructions. The labeled ILT7 was adjusted to 1µM with a buffer containing 20mM HEPES pH 7,5, 150mM NaCl and supplemented with 0.05 % Tween 20. A series of 16 1:1 dilutions of BST2 WT or mutants was prepared using the same buffer. For the measurement, each ligand dilution was mixed with one volume of labeled ILT7, and the samples were loaded into Monolith NT.115 Capillaries (NanoTemper Technologies). Instrument parameters were adjusted to 80% LED power and medium MST power (40%). Data of three independently pipetted measurements were analyzed (MO.Affinity Analysis software version 2.3, NanoTemper Technologies) using the signal from an MST-on time of 20s.

### Biophysical characterization of BST2 by circular dichroism (CD)

CD spectroscopy measurements were performed using a JASCO spectropolarimeter equipped with a thermoelectric temperature controller. Spectra of each sample were recorded at 20 °C in buffer A (50 mM phosphate pH 7.5). For thermal denaturation experiments, the ellipticity was recorded at 222 nm with 1 °C steps from 20 to 92 °C with a slope of 1°/min. Ellipticity values were converted to mean residue ellipticity.

### Crosslinking

BST2 WT and mutants were crosslinked with 5mM EGS (ethylene glycol bis succinimidylsuccinate, Pierce) at 1mg/ml in a buffer containing 50 mM phosphate pH 7.5 for 15 min at room temperature. Crossed linked samples were separated on a 15% SDS-PAGE and stained with Coomassie Brilliant Blue.

### Statistical analysis

Statistical analysis was performed using repeated measures ANOVA, with Bonferroni’s multiple comparison test or two-tailed paired Student’s t-tests. A p value of <0.05 was considered significant. The following symbols were used throughout the manuscript: *** p<0.001, ** p<0.01, * p<0.05, ns not significant (p>0.05).

## Acknowledgements

We thank E. Massicotte and J. Lord-Grignon for assistance with flow cytometry; the IRCM clinic staff, and all volunteers for providing blood samples. The plasmids encoding the BST2 ectodomain with the non-overlapping quadruple alanine mutations were a kind gift from Dr. Paul Spearman (University of Cincinnati). The ILT7 reporter cell lines were a generous gift from Dr. Y-J Liu.

This study was supported by Canadian Institutes of Health Research (CIHR) grants PJT148686 and FDN 154324 to ÉAC. ÉAC is recipient of the IRCM-UdeM Chair of Excellence in HIV research. WW acknowledges support from the ANRS (N° 073744), the Institute Universitaire de France (IUF) and the Grenoble Partnership for Structural Biology platforms (ISBG; UMS 3518 CNRS-CEA-UJF-EMBL) funded by FRISBI (ANR-10-INSB-05-02) and GRAL (ANR-10-LABX-49-01). NA was supported by a PhD fellowship from the Fond National de la Recherche Luxembourg. We further thank Caroline Mas (ISBG, Biophysical Platform Manager) and Pierre Soule (Application specialist, Nanotemper society) for their expert help with the analyses of the Microscale Thermophoresis experiments.

## Author Contributions

MGB and EAC conceived the original idea with input from WW. MGB, AL, and AM designed and carried out the functional experiments while NM, NA designed and carried out the binding experiments and the biophysical studies with input from FG and WW. MGB and EAC wrote the manuscript with support from WW, NM, and AL. EAC supervised the project with input from WW.

---

**Table S1.**
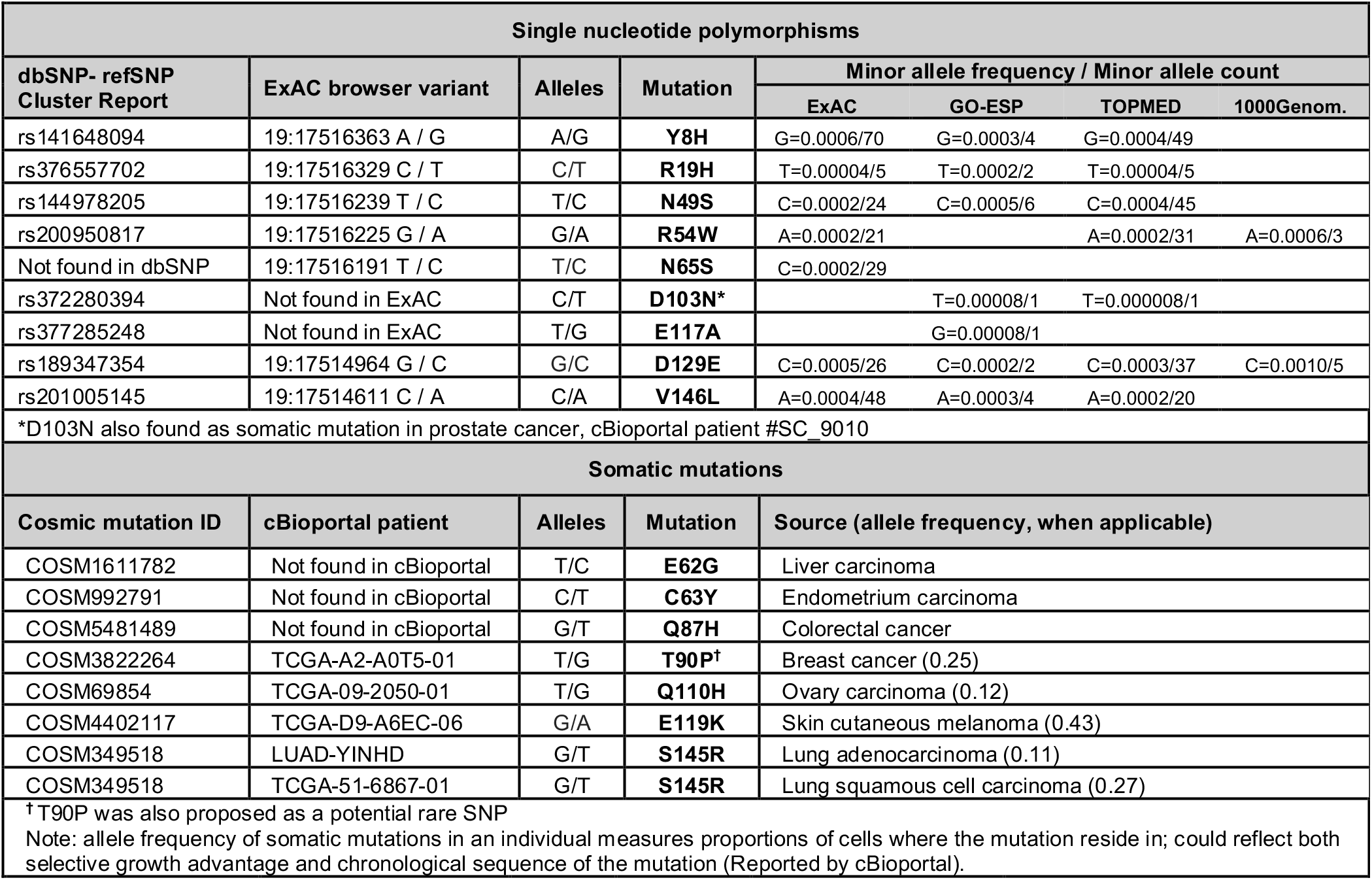
Natural variants of BST2

**Table S2.**
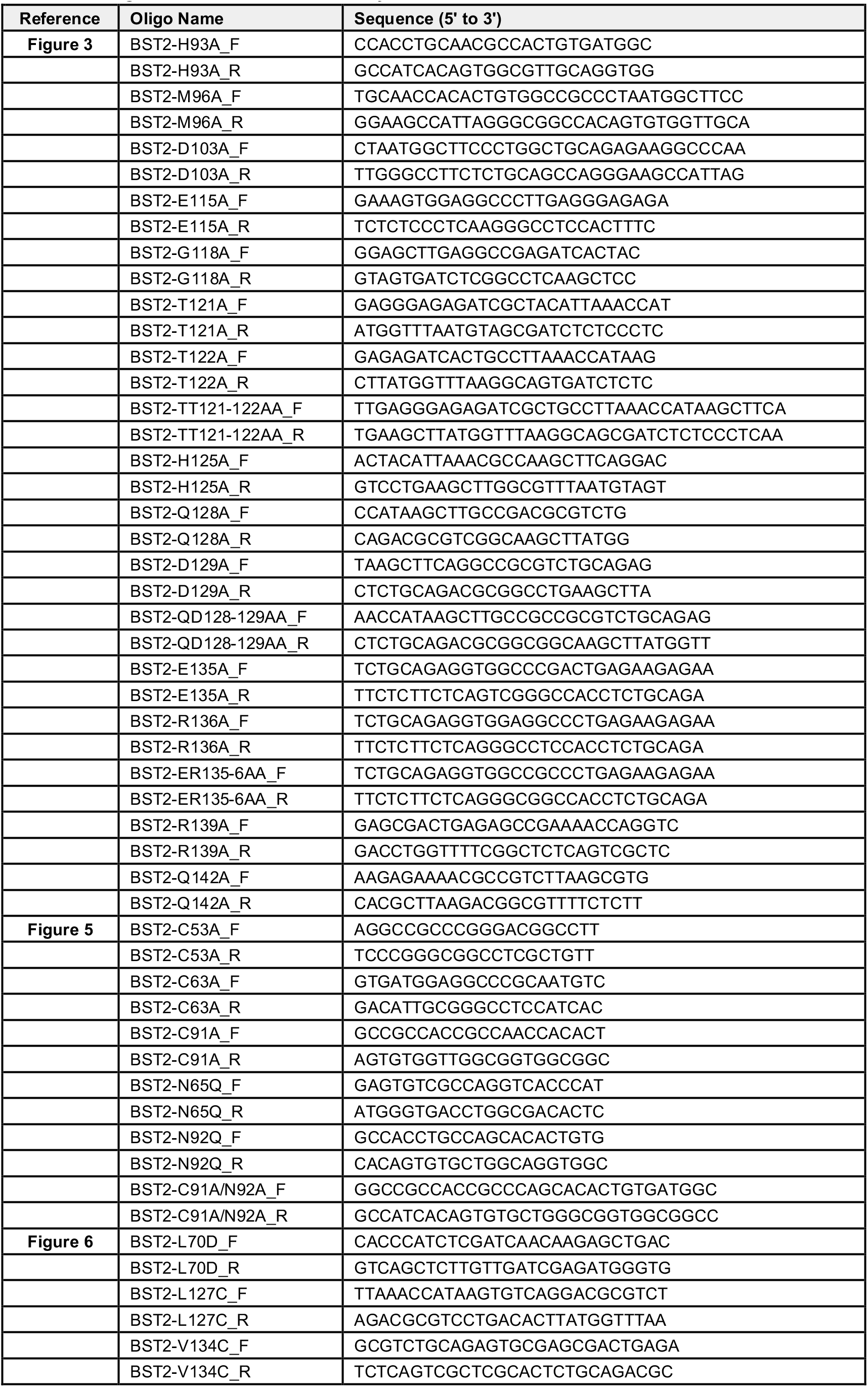
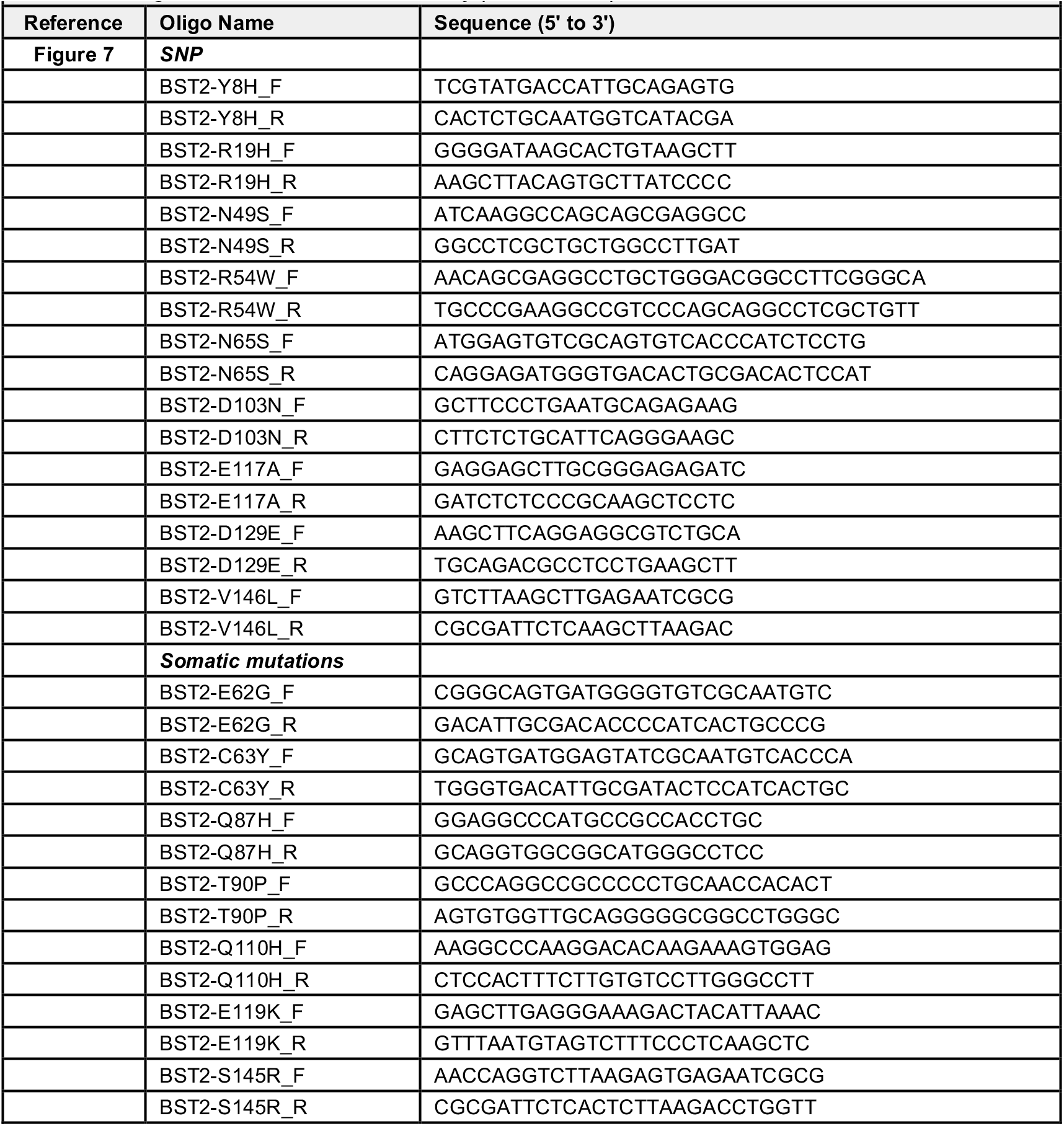
Oligonucleotides used in this study

