## Supplementary files for "ILT7 activation and plasmacytoid dendritic cell response are governed by BST2 determinants that are structurally-distinct"

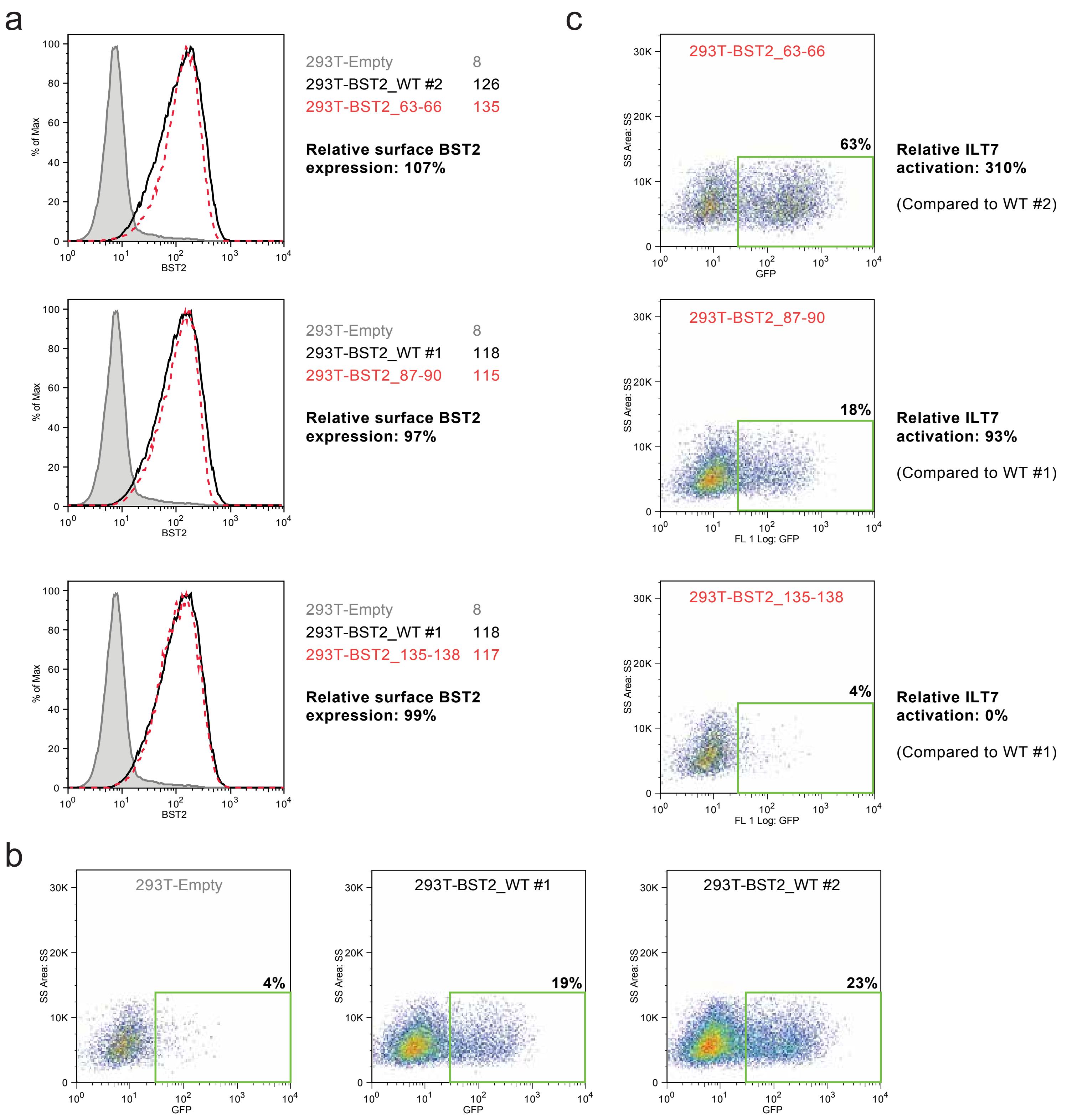


**Fig S1- Standardization of ILT7+ NFAT-GFP reporter assays. a,** Representative examples of flow cytometry analysis of surface BST2 in HEK-293T cells after transfection of control (293T-empty, solid grey histogram) or the indicated BST2 plasmids (black solid lines for BST2 WT or red dashed lines for BST2 mutants), 48 hours post transfection (hpt). Mean fluorescence intensity (MFI) values are indicated for each sample. Relative surface BST2 expression for each mutant compared to the corresponding BST2 WT control are indicated. In the examples shown, two different concentrations of BST2 WT were used (#1 and #2) in order to better match the surface expression levels of the different mutants tested. (B-C) ILT7+ NFAT-GFP reporter cells were co-cultured with HEK-293T (empty control) or the BST2-expressing HEK-293T cells for an additional 18-24 h. Co-cultures of transfected HEK-293T cells and ILT7+ NFAT-GFP reporter cells were then stained for ILT7 and analyzed by flow cytometry. Reporter cells were identified as ILT7-positive, and the percentage of GFP-positive cells in this population was calculated. **b,** Representative example of ILT7 activation as determined by the percentage of GFP-positive cells (indicated for each sample) measured for negative (empty) or positive controls (BST2 WT #1 and #2) co-cultures. **c,** Representative example of ILT7 activation as determined by the percentage of GFP-positive cells (indicated for each sample) measured for co-cultures between ILT7+ NFAT-GFP reporter cells and HEK-293T cells expressing specific BST2 mutants from panel (A). For each mutant the relative ILT7 activation (compared to its respective BST2 WT control) is also indicated.

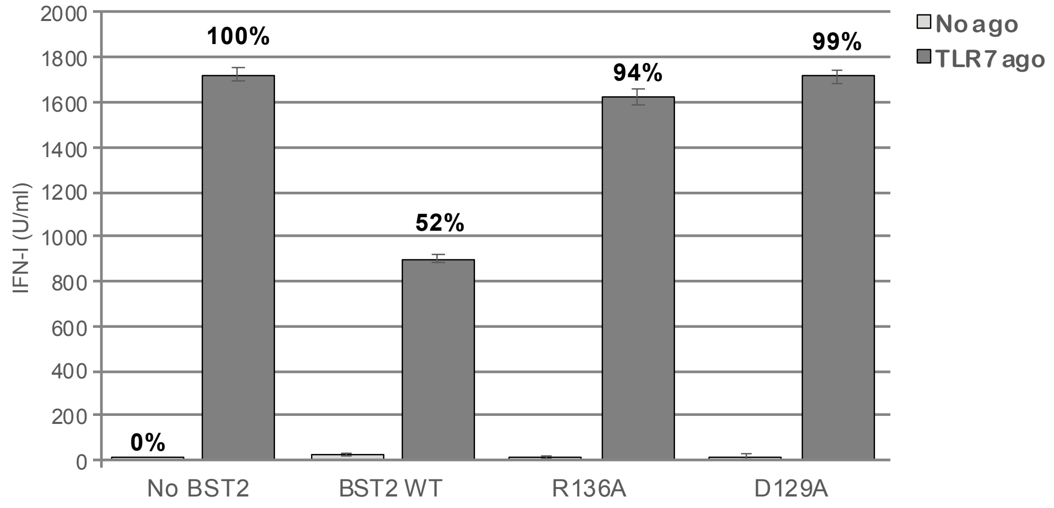


**Fig S2- Standardization of PBMC-based assays.** HEK-293T cells were transfected with either an empty plasmid (no BST2) or plasmids expressing BST2 WT 48 h prior to co-culture with PBMCs. After 4 h of co-culture, samples are either untreated or treated with Gardiquimod (TLR7 agonist) and levels of bioactive IFN-I released in supernatants measured 18-24 h later, as described in the Methods section. Representative examples of the absolute levels of IFN-I released (U/ml) and relative percentage of IFN-I released by PBMCs in contact with HEK-293T (no BST2) or BST2-expressing HEK-293T (WT or mutants) in the presence or the absence of TLR7 agonist are shown. The level of IFN-I detected in the co-culture of PBMC and HEK-293T (no BST2) in the presence of TLR7 agonist was set at 100%.

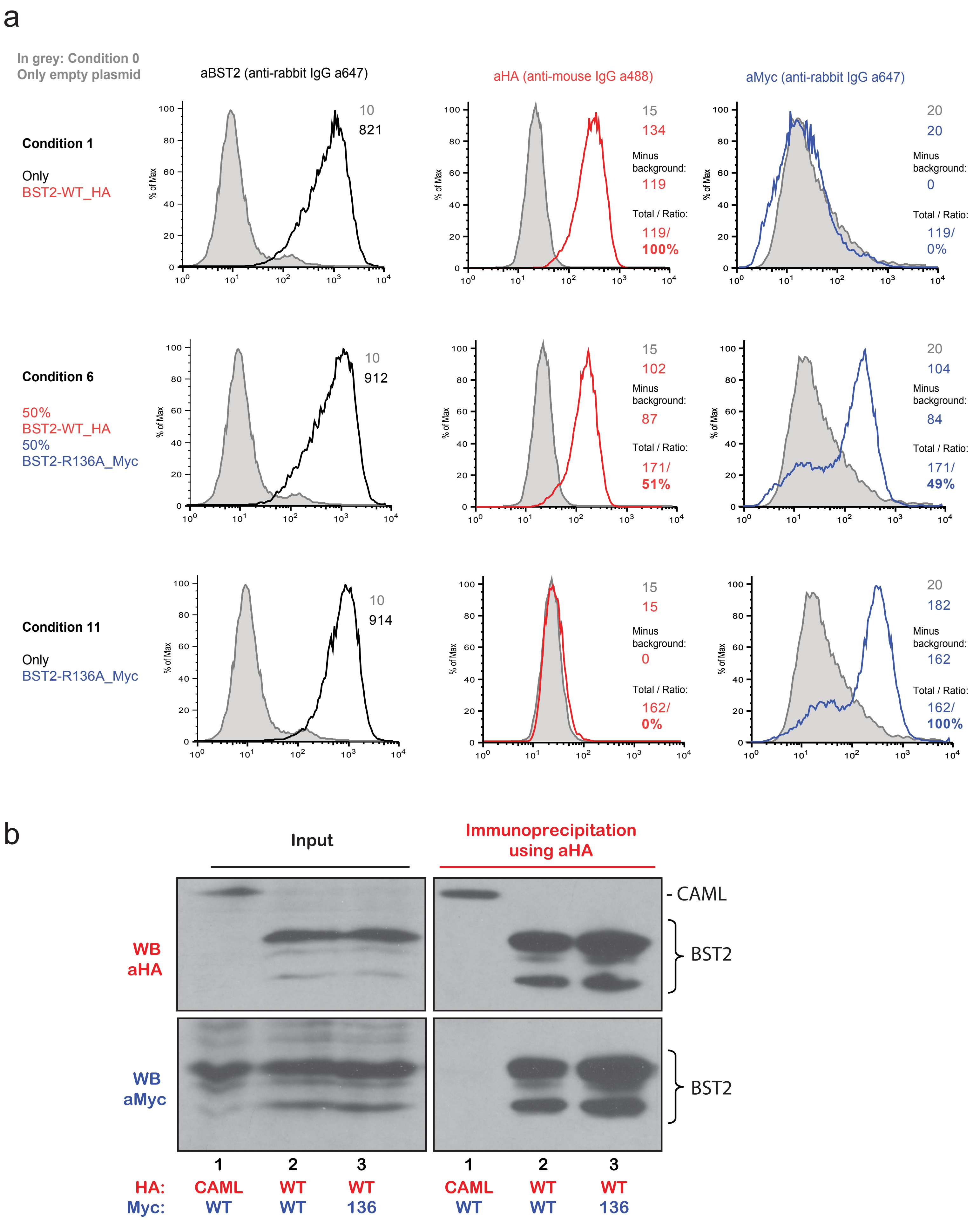
a
In grey: Condition 0
Only empty plasmid
aBST2 (anti-rabbit IgG a647)
aHA (anti-mouse IgG a488)
aMyc (anti-rabbit IgG a647)
Condition 1
Only
BST2-WT_HA
10
821
15
134
Minus background:
119
Total / Ratio:
119/
100%
20
20
Minus background:
0
Total / Ratio:
119/
0%
Condition 6
50%
BST2-WT_HA
50%
BST2-R136A_Myc
10
912
15
102
Minus background:
87
Total / Ratio:
171/
51%
20
104
Minus background:
84
Total / Ratio:
171/
49%
Condition 11
Only
BST2-R136A_Myc
10
914
15
15
Minus background:
0
Total / Ratio:
162/
0%
20
182
Minus background:
162
Total / Ratio:
162/
100%
% of Max
b
Input
Immunoprecipitation using aHA
- CAML
BST2
BST2
WB aHA
WB aMyc
1 2 3
1 2 3
HA: CAML WT WT
Myc: WT WT 136
CAML WT WT
WT WT 136

**Fig S3- Standardization and controls for heterodimerization assays. a,** Surface expression of HA-BST2 WT and Myc-BST2 R136A in transfected HEK-293T cells as determined by flow cytometry. Cells were stained with anti-HA (red histogram), anti-Myc (blue histogram), or anti-BST2 (black histograms- for total BST2) antibodies. Staining for total BST2 was used as an internal control to ensure that similar levels of surface BST2 were achieved in all conditions. Representative staining examples are shown for condition 0 (cells transfected with empty plasmid, grey solid histograms), conditions 1 and 11 (cells that received only HA-BST2 WT or Myc-BST2 R136A plasmids, respectively) and condition 6 (equimolar ratios of each plasmid). MFI values are indicated for each condition on the top (black for anti-BST2 staining, red for anti-HA staining, blue for anti-Myc staining and grey for negative control cell staining for all antibodies used). For each condition, MFI of negative control was subtracted from anti-HA and from anti-Myc MFI values (value indicated as Minus background) and the subtracted MFI values were added to generate a Total value. Finally, the Total and subtracted MFI values were used to generate the ratios of each isoform (expressed as % of Total). These calculated ratios were plotted for conditions 1 to 11 in Fig. 6A. **b,** Co-immunoprecipitation analysis of protein lysates of HEK-293T cells expressing either HA-CAML (used as a membrane-associated protein control) and Myc-BST2 WT (lanes 1) or HA-BST2 WT and Myc-BST2 WT (lanes 2), or HA-BST2 WT and Myc-BST2 R136A (136) (lanes 3). Immunoprecipitation was performed using mouse anti-HA mAb and the immunocomplexes were further analyzed by western blot using mouse anti-HA mAb or rabbit anti-Myc polyclonal antibodies, as indicated.

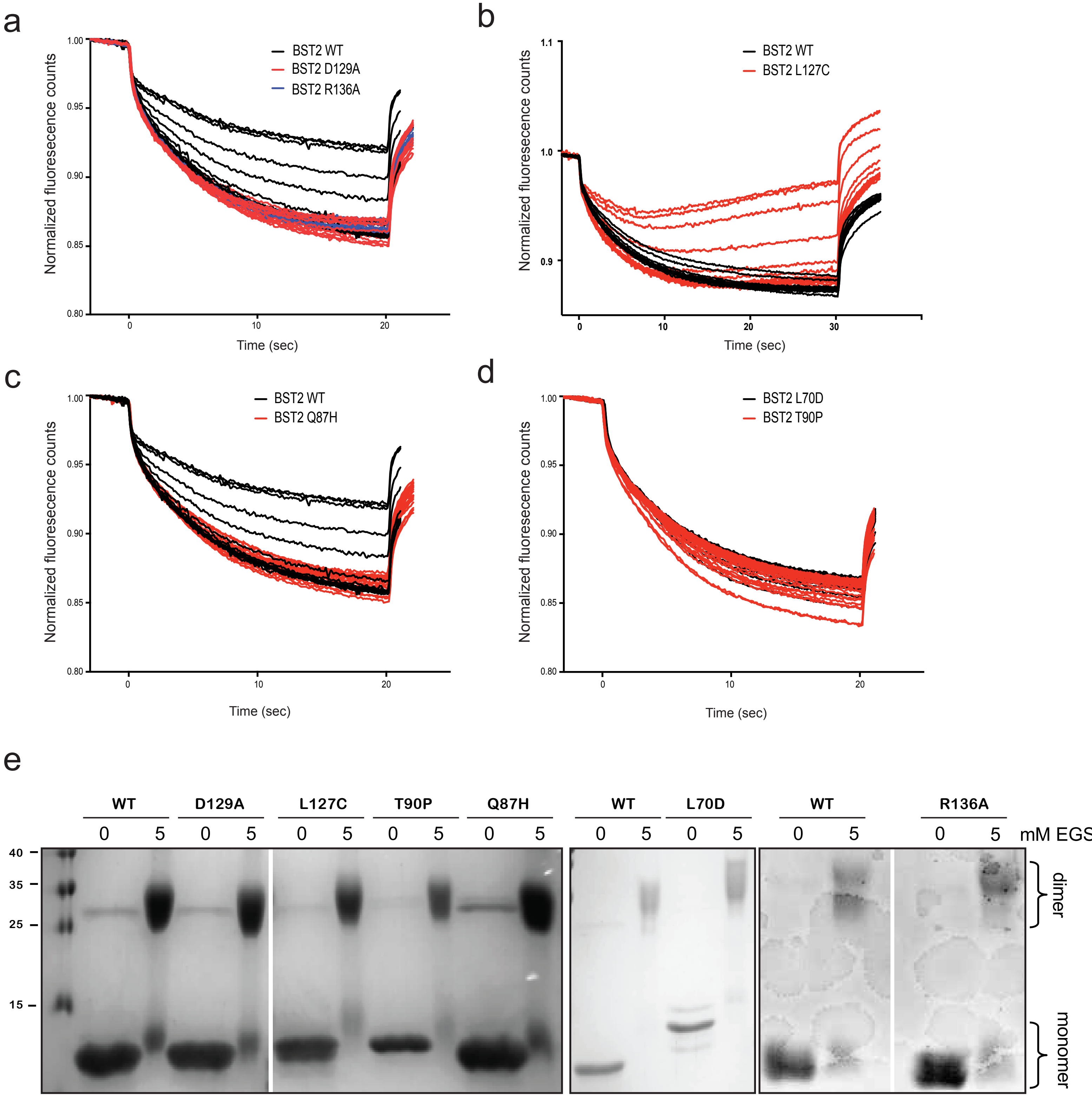


**Fig S4- Thermophoretic analysis of BST2–ILT7 interaction: microscale thermophoresis (MST) traces.** Recombinant GST-BST2 (47-159) WT or mutants were mixed at different concentrations with fixed amounts of bacILT7 24-223. MST traces of ILT7 towards **a,** BST2 WT (black) or mutants D129A (red) and R136A (blue), **b,** BST2 WT (black) or mutant L127C (red), **c,** BST2 WT (black) or mutant Q87H (red), and **d,** mutants L70D (black) and T90P (red) were generated. All experiments were done in triplicates. **e,** BST2 WT and mutants (1mg/ml) were cross-linked with 5mM EGS for 15 min at room temperature. Crossed linked samples were separated on a 15% SDS-PAGE and stained with Coomassie Brilliant Blue. Regions corresponding to monomeric and dimeric forms of BST2 are shown. Each mutant was paired with one BST2 WT analyzed from the same SDS-PAGE (samples corresponding to the same SDS-PAGE were boxed).

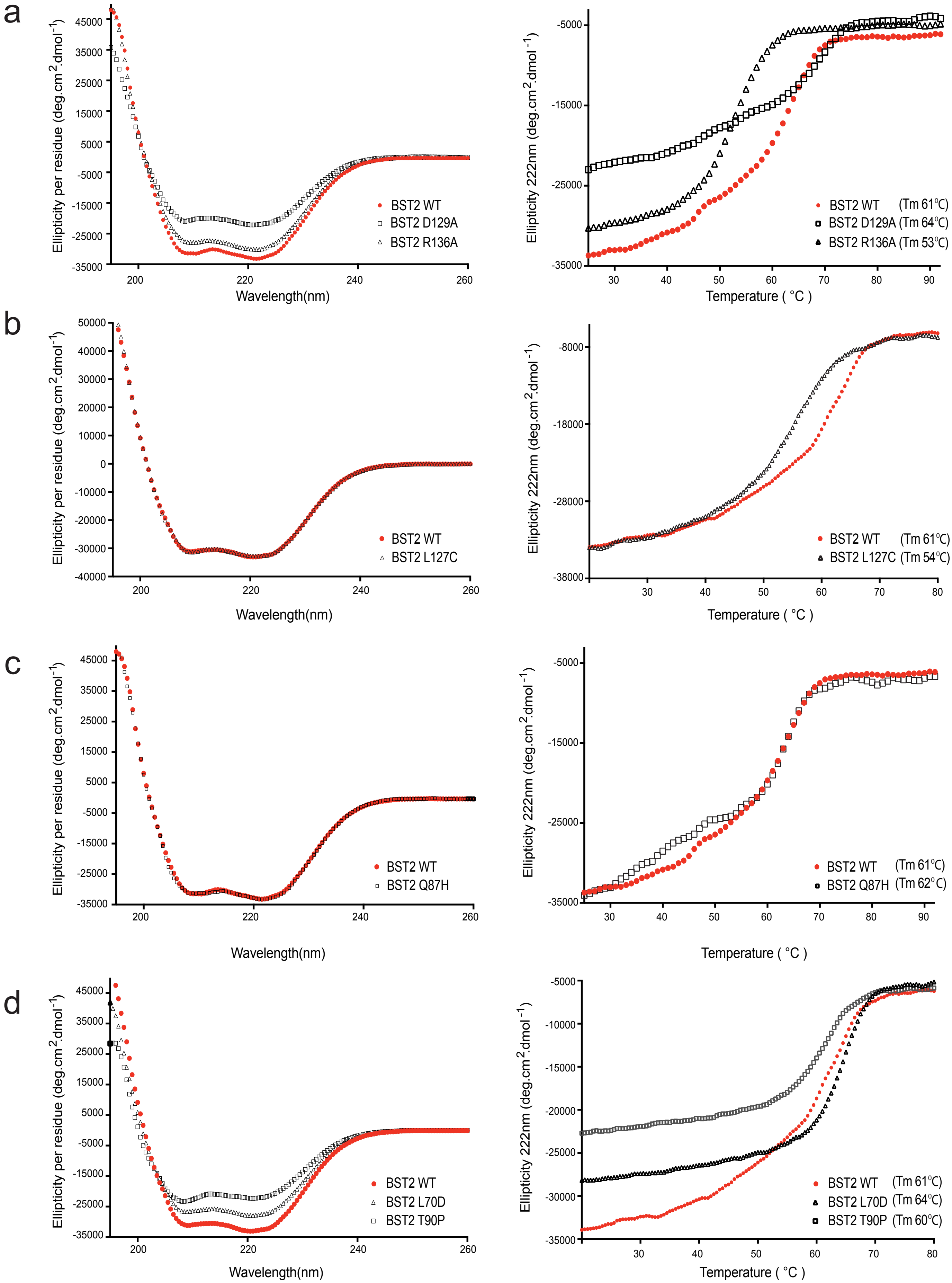


**Fig S5- Circular Dichroism and thermal denaturation studies.** Circular dichroism spectra were recorded from 260 to 190 nm (left panels). Thermostability measurements were performed at 222 nm (right panels) for recombinant GST-BST2 (47-159) WT (red circles) and respective BST2 mutants, **a,** D129A (back squares) and R136A (black triangles), **b,** L127C (black triangles), **c,** Q87H (back squares), and **d,** L70D (black triangles) and T90P (back squares). Melting temperatures (Tm) were extrapolated form the curves and indicated in parenthesis next to each sample.
